# Postprocessing-Enhanced Machine Learning for Reliable Real-Time Sleep Staging in Closed-Loop Neuromodulation

**DOI:** 10.1101/2025.09.22.677765

**Authors:** Tony Sungjin Chae, Jinmo Jeong, Kevin Kai Wing Tang, William D. Moscoso-Barrera, Benjamin Baird, Xue-Xin Wei, José del R. Millán, Huiliang Wang

**Author notes:** Author to whom any correspondence should be addressed.

## Abstract

Real-time sleep stage classification is important for closed-loop neuromodulation at certain stages during sleep, yet current models often yield noisy and unstable outputs that risk false triggers. These fluctuations, especially near stage boundaries, can compromise the safety and reliability of stimulation. Existing methods frequently rely on model-specific architectures or require extensive tuning to maximize precision, which limits their generalizability and hinders deployment in real-time systems. To address this, we propose a lightweight, classifier-independent post-processing pipeline that stabilizes predictions without modifying the underlying classifier. Our method first applies temporal smoothing to predicted sleep-stage probabilities, followed by conservative control logic to enhance stimulation precision. We evaluate four smoothing techniques: Moving Average (MA), Exponential Smoothing (ES), Kalman Filtering (KF), and a novel Weighted Exponential Smoothing (WES), on a 29-subject open-source sleep dataset. To capture the trade-offs between stability and responsiveness, we introduce new real-time evaluation metrics. We identify an optimal smoothing intensity range and assess three control strategies: probability thresholding, naïve waiting, and entropy constraint, as well as a hybrid method combining high confidence and low uncertainty. Smoothing improves generalization and robustness: under both low and high synthetic noise (*σ >* 0.2), smoothed outputs retain 10–15% higher precision and recall across all stages. Control logic further enhances reliability: our hybrid method achieves 80% (Wake), 96% (N2), 98% (N3), and 86% (REM) precision. Finally, we introduce a stage-transition constraint matrix to suppress biologically implausible transitions (e.g., REM→ N3), to further stabilize outputs during evidence accumulation, with potential applications in objective sleep quality assessment. To our knowledge, this is the first study to systematically characterize real-time trade-offs between smoothing, latency, and control logic in sleep staging. Overall, our generalizable framework is expected to improve safety, interpretability, and deployment feasibility of real-time sleep classification for both clinical and wearable closed-loop neuromodulation systems.

## 1. Introduction

Accurate real-time sleep stage classification is a key enabling technology for closed-loop neuromodulation therapies and sleep research interventions. Prior studies have demonstrated that real-time closed-loop acoustic stimulation can reduce sleep onset latency [1, 2]. Similarly, adaptive deep brain stimulation (DBS) has been applied in Parkinson’s disease patients to selectively target N3 sleep, enabling stage-specific therapeutic effects [3]. Non-invasive approaches such as transcranial magnetic stimulation (TMS) have also been explored to enhance slow-wave sleep or facilitate memory consolidation during NREM stages [4]. Notably, the amplification of sleep spindles via transcranial alternating or direct current stimulation (tACS, tDCS) has shown potential in modulating sleep architecture [5, 6]. These applications all rely on the system’s ability to accurately detect transitions into specific sleep stages and to deliver stimulation at precisely the right moment, despite the subject being unconscious and unable to provide feedback. This demands a high level of reliability and stability in automated sleep staging [7, 8].

Over the past decade, significant progress has been made in offline automatic sleep staging using machine learning and deep neural networks, achieving high accuracies on benchmark datasets. However, deploying these classifiers in an online, closed-loop setting still presents key challenges in accurately determining stimulation timing and delivering precise feedback in real time [7, 9]. Existing clinical approaches often rely on model-specific parameters or hand-tuned delays to increase precision, which require extensive tuning across subjects and do not generalize well to new models or datasets. Moreover, due to their complexity, many high-performing offline models are not directly applicable to online systems [10, 11]. For instance, in a system explicitly designed for real-time stimulation, Nguyen et al. reported frequent signal degradation that necessitated offline smoothing to recover unscored epochs, highlighting the vulnerability of online staging under real-world conditions [1].

Real-time sleep stage classification is inherently susceptible to noisy and jittery outputs, as both internal and external noise sources can induce momentary misclassifications or stage oscillations —even in models with high average accuracy [10]. While such fluctuations may be inconsequential in offline analyses, they pose critical challenges for live, closed-loop interventions. This vulnerability is especially exposed in sleep, where subjects are unconscious and have an absence of volitional control, unlike typical BCI (Brain-Computer Interface) systems that operate under user intention.

The sources of instability can be broadly categorized into three types. (1) Mechanical noise, including electrode drift, electromagnetic interference (EMI), power line noise [12], and stimulation-induced artifacts [13] such as encapsulation effects. (2) Biological noise, arising from non-stationary EEG (electroencephalogram) patterns with overlapping spectral features, EMG (electromyography) bursts, and ambiguous transitions between brain states that result in fuzzy or dynamic decision boundaries that further complicate the interpretability of the underlying signal [14–16]. (3) Model and data noise, which stems from limitations in both training data and learning algorithms. These include small training sample sizes, subject variability, labeling inconsistency, classifier uncertainty, weak generalization, and noisy feature extraction from short temporal windows [17, 18]. These multifactorial noise sources amplify uncertainty and false positives in real-time sleep stage classification models and underscore the need for robust temporal stabilization strategies.

False-positive triggers not only will reduce the efficacy of the intervention (by delivering stimuli at the wrong times or too often) but could also disrupt sleep and pose safety concerns [8]. Therefore, achieving stable, high-precision sleep stage detection in real time is both crucial and challenging. Sleep stage detection is a classification problem, where predicted probabilities (e.g., softmax outputs [19]) are often miscalibrated and may not reflect true correctness likelihoods due to output instability. One way to improve the reliability of stimulation systems is to redistribute and constrain the classifier’s posterior probabilities into a new temporal series *P*_*t*_, which functions as a low-pass filter to enforce smoothness and robustness in the output. Post-processing techniques like temperature scaling [20] help recalibrate these naïve confidences, while temporal smoothing further reduces sequential noise, yielding more trustworthy predictions over time. In BCI research, for example, evidence accumulation techniques are used to improve the reliability of asynchronous control signals. Millán et al. demonstrated that applying a smoothing filter over a motor imagery classifier’s output eliminated the uncertainty of single-sample EEG predictions and significantly reduced random false positives during idle periods [21]. Their implementation of a leaky integrator (exponential smoothing) or more sophisticated Naïve Bayesian integration yielded much steadier control without sacrificing responsiveness. Inspired by this, one can posit that temporal smoothing might similarly benefit sleep stage classifiers, which also operate on a sequential data stream and face similar issues of uncertain, oscillatory predictions. Indeed, preliminary work in the sleep domain hints at the value of smoothing: Wang et al. proposed an “automatic sleep level” for nap monitoring based on the conditional probability of stages, smoothed over time, and found it useful with nap sleep physiology [22]. To our knowledge, no prior study has systematically defined real-time smoothing metrics or compared different smoothing strategies for sleep staging in a closed-loop context. Although post-processing methods exist, there is little guidance on selecting appropriate strategies for different real-time applications, or how their trade-offs quantitatively compare.

Our aim is therefore: (1) to develop a classifier-independent framework that improves model performance, stability and interpretability in real-time sleep stage classification, and (2) to systematically compare smoothing and control strategies in order to guide optimization and maximize precision, particularly in unseen subjects and practical closed-loop settings.

To address this gap, we systematically designed two categories of post-processing: (1) temporal smoothing and (2) conservative control logic. These modules were implemented after the classifier in a real-time closed-loop sleep modulation pipeline, following standard preprocessing steps such as artifact rejection, Independent Component Analysis (ICA), filtering, and feature extraction. End-to-end system pipeline is described in Fig. 1. Since our method operates directly on softmax-based posterior probabilities, it is broadly compatible with any classifier architecture that outputs class probability distributions, including traditional neural networks [23], graphical models [24], convolutional neural networks (CNNs) [25], hybrids of CNNs and long short-term memory networks (LSTMs) [26, 27], and Transformer-based architectures [28, 29].

**Figure 1.**
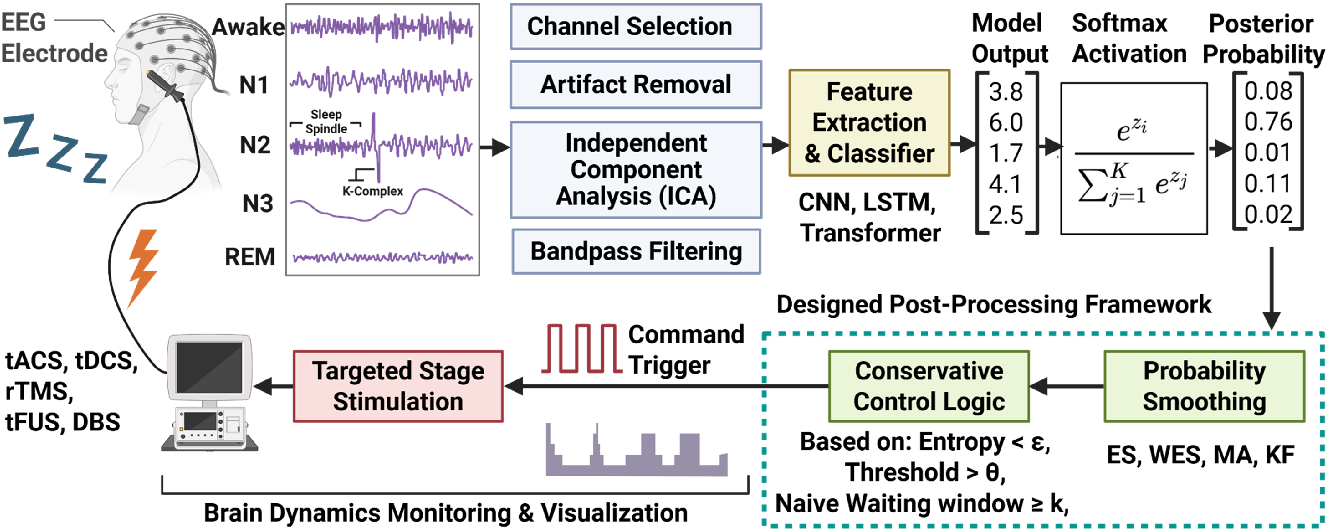
Designed Real-time Closed-loop Sleep Modulation Pipeline for Increased Precision and Medical Safety. In the traditional closed-loop stimulation paradigm for awake/sleep (N1-N3, REM) neuromodulation, the decoder records neurophysiological signals through electrodes, preprocess the data (e.g., channel selection, artifact removal, Independent Component Analysis, band-pass and notch Filtering), extracts features and trains on those signals, then classifies the current sleep stage based on the posterior probability output of the model; The Softmax function transforms a vector of numerical values of model output into a probability distribution, where each element in the output vector represents the probability of belonging to a specific class. Since our smoothing framework operates on softmax-based posterior probabilities, it is compatible with any architecture that outputs class distributions, without modifying the classifier’s internal structure or training. This makes the method broadly applicable: whether one uses a simple feature-based neural network (as in our case) or a state-of-the-art deep learning model. This includes ANNs [23], CNNs [25], LSTMs [26], CNN-LSTM hybrids (e.g., DeepSleepNet [27]), graph-based models [24], and Transformers [28, 29]. Typically, when the target stage is detected, the command trigger is sent to the stimulation device (e.g., tACS, tDCS, rTMS, tFUS, DBS) and the stimulus is delivered to the subject. Brain dynamics are monitored and visualized throughout the stimulation process. In out study, real–time–compatible post-processing stages (green) are added after raw model output to stabilize sleep-stage estimates while preserving rapid response to genuine transitions, suppress false positives, and yield more reliable control logic for targeted stimulation. This includes two steps: Probability smoothing and Conservative control logic. Exponential Smoothing (ES), Weighted Exponential Smoothing (WES), Moving Average (MA), and Kalman Filtering (KF) were implemented and tested to compare probability stability (entropy), delay, mismatch rates, and overall model performance on raw versus smoothed outputs. WES and Transition-Constraint WES (TC-WES) were benchmarked against traditional smoothing (ES, MA, KF) to assess whether non-recursive evidence accumulation with/without neurophysiological constraints enhances sleep-stage classification without retraining the base decoder. Various conservative control logics (threshold-cutoff, naive waiting, entropy-constrained, hybrid constraint (threshold+entropy) were compared to maximize precision and manage the precision–recall trade-off while avoiding false positives and to maximize precision. This classifier-independent post-processing method could be applied to any given sleep staging model that generates a posterior probability, making this approach a generalizable and transferable pipeline for real-time sleep neuromodulation.

First, we evaluated four smoothing methods to assess their impact on model performance and transition stability, delay, and entropy, to identify optimal smoothing intensity. We also explored physiologically implausible sleep stage sequences into an evidence accumulation process to reduce false positives, extending prior work that used discrete Hidden Markov Models (DHMM) to constrain stage transitions [30], evaluating its applicability to neural network–based models.

Second, we examined three control logics to evaluate whether they could enhance stimulation precision when applied after smoothing by constraining the model output.

While prior studies have relied on threshold-based and fixed-delay strategies to improve precision [31–35], these methods often trade off recall and responsiveness: thresholds are vulnerable to noise, and rigid delays risk missing brief but clinically relevant windows, particularly in short or phasic stages like N1 or REM [36]. Despite their widespread use, these control strategies have never been systematically compared under a unified framework, leaving practitioners without clear standards for selecting methods suited to specific real-time applications. Our standardized evaluation would not only simplify the optimization process but also allow users to flexibly choose control logic based on application-specific priorities, such as minimizing false positives or ensuring rapid response to brief stages. Together, these components form a unified, classifier-independent, real-time post-processing framework (Fig. 1, green blocks), designed to stabilize sleep-stage classification and improve reliability in closed-loop neuromodulation systems.

## 2. Methods

### 2.1. Dataset and Preprocessing

We use the publicly available Open Source Framework for Analyzing Physiology (OSF-ANPHY) sleep dataset, which comprises overnight high-density EEG polysomnography recordings from 29 healthy adults [37]. Each recording included the full standard sleep monitoring channels: EEG, electrooculogram (EOG), and EMG. The dataset included high-density EEG channels (a total of 83 channels) and accompanying expert-scored sleep stage annotations for every 30-second epoch (window), following the American Academy of Sleep Medicine (AASM) standard [38]. For our analysis, we focused on a subset of signals suitable for real-time wearable device implementation. Specifically, we selected 6 channels consisting of 4 EEG (including a non-traditional frontal-lateral montage electrode to capture frontal slow waves and localized activity) channels of T3-Ref (T3), F7-Ref (F7), Fp1-Ref (Fp1), FPz-Ref (FPz), 1 EOG channel of LOC(left outer canthus)-Ref, and 1 EMG channel of left-chin. Our channel selection is consistent with ongoing neuromodulation studies that commonly target frontal and temporal regions (e.g., Fz, Fp1, Fp2, F7, T3) to modulate and monitor slow oscillations and spindles during sleep [39–41]. Additionally, we adopt a minimal yet spatially distributed montage to cover anterior to posterior regions, aligned with recent closed-loop wearable EEG systems that demonstrated reliable sleep feature detection using reduced channels [1, 42].

This channel configuration captures the core physiological signatures used in manual staging: EEG oscillations, eye movements, and muscle tone, while limiting data volume for faster processing. All signals were sampled at 512 Hz and notched-filtered (59-61 Hz) to remove power line noise. Next, we applied a third-order Butterworth band-pass filter in the typical sleep frequency range (0.5–45 Hz for EEG, 0.1-35 Hz for EOG, and 0.1-150 Hz for EMG) to remove drift and high-frequency noise. Lastly, we applied Independent Component Analysis (ICA) to remove EOG and EMG artifacts in EEG signals. Each 30-second window was then segmented for feature extraction. We partitioned the 29 subjects into a training set (20 subjects), a validation set (5 subjects), and a hold-out test set (4 subjects). This leave-subjects-out split ensured that the final evaluation tested generalization to entirely unseen individuals – a critical requirement for practical sleep staging models. The validation set was used to evaluate trade-offs between performance metrics, optimize smoothing method hyperparameters, and decision thresholds. All final results are reported on the 4 test subjects.

### 2.2. Feature Extraction and Model Construction

#### Model Construction

We implemented a lightweight feature-based deep neural network classifier suitable for real-time deployment (e.g. on wearable devices or bedside processors). From each 30 s epoch of 6-channel data, we extracted a total of 118 scalar features capturing time domain and frequency-domain characteristics known to correlate with sleep stages. For the EEG channels, these included: bandpower features (relative and absolute power in alpha, beta, theta, delta, gamma, spindle frequency bands), burst/suppression metrics, and slow-wave characteristics; for the EOG: energy in low-frequency bands detection features; and for the EMG: average amplitude and variance (as indicators of muscle tone) among others. More details can be found in Feature Extraction section below. All features were then standardized (z-scored) across the training set. Using these features, we trained a fully connected artificial neural network (ANN) classifier with 4 hidden layer (sizes 512, 512, 256, and 128 neurons respectively [43], LeakyReLU activation [44], 0.5 dropout was applied [45]) and an output softmax layer [19] yielding posterior probabilities for each of the 5 sleep stages (Wake, N1, N2, N3, REM). The network was trained on the 20-subject training set with cross-entropy loss with class weights and early stopping. The Adam optimizer for learning rate of 1e-4 and weight decay of 1e-5 was used. Despite its simplicity [46], this ANN achieved high performance with typical deep learning models on similar data. The 29-group K-fold was used to avoid subject data mixture while validating baseline model performance. (Detailed baseline model performance given in Results). Importantly, the model outputs a probability distribution over stages for each epoch, which we leverage in the smoothing step. By using a small, feed-forward architecture and a limited feature set, we ensured the classifier’s computations per epoch were minimal (on the order of milliseconds on a standard CPU), making it feasible for real-time operation (30 s latency for the epoch itself, plus negligible computation overhead).

#### Feature Extraction

We extracted both frequency-and time-domain features from 30-second windows of EEG, EOG, and EMG signals. Features were computed per channel using Welch’s method for power spectral density (PSD) estimation and basic statistical operations in the time domain [47]. The sampling frequency was set to 512 Hz. Table. 1 lists all types of feature used in this paper to construct the baseline model.

**Table 1.** Extracted Features from EEG, EOG, and EMG Channels.

| No. | Type | Feature Description | Source |
| --- | --- | --- | --- |
| 1 | PS | Total power of 0.1–30 Hz | EEG |
| 2 | PS | Total power of 0.1–30 Hz | EMG |
| 3 | PS | Total power of 0.1–30 Hz | EOG |
| 4 | PR | Delta ratio (0.1–4 Hz / total) | EEG |
| 5 | PR | Theta ratio (4–8 Hz / total) | EEG |
| 6 | PR | Alpha ratio (8–13 Hz / total) | EEG |
| 7 | PR | Beta ratio (22–30 Hz / total) | EEG |
| 8 | PR | Gamma ratio (30–45 Hz / total) | EEG |
| 9 | PR | Spindle ratio (11–16 Hz / total) | EEG |
| 10 | PR | Delta-to-Alpha Ratio (DAR) | EEG |
| 11 | SF | Mean frequency of 0.1–30 Hz | EEG |
| 12 | SF | Mean frequency of 0.1–30 Hz | EMG |
| 13 | SF | Mean frequency of 0.1–30 Hz | EOG |
| 14 | DR | Spindle duration ratio (sec / 30s) | EEG |
| 15 | PR | Slow eye movement ratio (0.1–4 Hz / total) | EOG |
| 16 | TD | Time-domain mean | EEG, EOG, EMG |
| 17 | TD | Time-domain variance | EEG, EOG, EMG |
| 18 | TD | Zero crossing rate | EEG, EOG, EMG |
| 19 | TD | Absolute mean amplitude | EEG, EOG, EMG |

#### 2.2.1. Frequency-Domain Features

##### Power Spectrum (PS)

We used Welch’s method with 4-second windows and 50% overlap to compute the PSD *P* (*f* ) of each channel. The total power in each band was computed as:

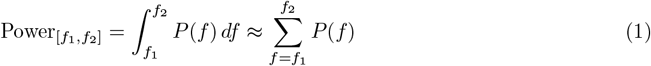

We defined the following frequency bands:

- Total: 0.1–30 Hz
- Delta: 0.1–4 Hz, Theta: 4–8 Hz, Alpha: 8–13 Hz
- Beta: 22–30 Hz, Gamma: 30–45 Hz, Spindle: 11–16 Hz

##### Power Ratios (PR)

Each band’s relative power was computed with respect to the total power:

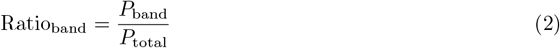

##### Delta-to-Alpha Ratio (DAR)

DAR is additionally implemented due to its proven benefits in sleep staging [48].

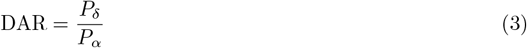

##### Slow Eye Movement Ratio (SR)

To characterize slow eye movements, we computed the slow ratio from the EOG channel:

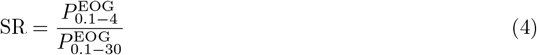

##### Spectral Frequency (SF)

For each band, the mean spectral frequency was computed as:

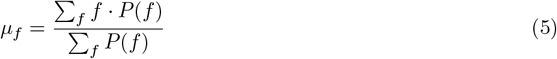

#### 2.2.2. Time-Domain Features

Time-domain statistics were computed on the raw signal within each 30s window:

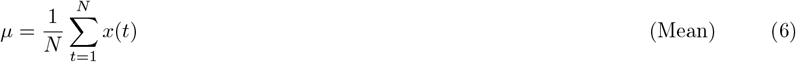

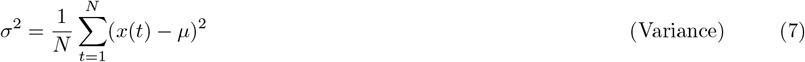

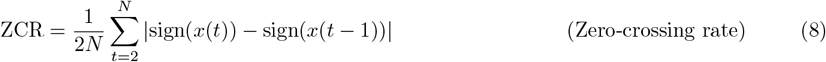

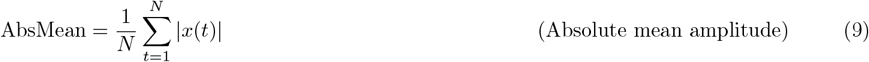

#### 2.2.3. Spindle Detection and Duration Ratio

We also added the spindle duration ratio and binary flag for additional features for N2 stage classification. Sleep spindles were detected based on criteria adapted from [49], using the following parameters: freq_sp_ = (12, 15) Hz, freq_broad_ = (1, 30) Hz, duration between 0.5–2s, min distance = 500 ms, and threshold settings corr ≥0.65, rel pow ≥0.2, RMS ≥1.5. A binary spindle flag was assigned if at least one spindle was detected in the 30s window. The spindle duration ratio (SDR) was computed as:

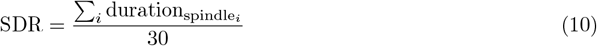

These features were extracted across all EEG, EOG, and EMG channels.

#### 2.3. Smoothing Method to posteriors

The first objective of our work was to quantify how various smoothing methods affect classification performance and temporal stability, by identifying optimal smoothing parameters and generalizability under practical constraints. Therefore, we selected four smoothing methods suitable for real-time systems: exponential smoothing [22, 50], moving average (commonly used in time-series signal processing [51]), Kalman filtering (widely applied in time-series tracking and control systems such as GPS [52,53]), and our proposed Weighted Evidence Smoothing (WES), a non-recursive approach to evidence accumulation. We additionally examined whether incorporating physiological transition constraints could further improve WES (TC-WES) by enforcing biologically plausible state progressions during the accumulation process.

Each method was applied to the sequence of softmax probability vectors *P*_*t*_ ∈ ℝ*C*, where *C* is the number of classes, to generate a smoothed estimate 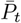. The final predicted class label *ŷ*_*t*_ is then obtained by:

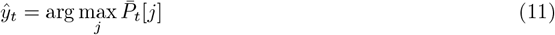

#### Exponential Smoothing (ES)

This method performs a weighted average recursively between the current prediction and the previous smoothed value:

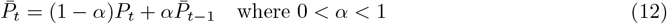

where *α* is the smoothing parameter.

#### Weighted Exponential Smoothing (WES)

To improve temporal stability in real-time sleep stage classification, we implemented a *Weighted Exponential Smoothing* (WES) approach to acquire more flexibility. This was inspired by ideas of evidence accumulation in probabilistic smoothing frameworks such as Naive Bayesian Integration [21]. Unlike traditional *Exponential Smoothing* (ES) that only compares previous window recursively, WES explicitly retains and averages past posterior outputs over a fixed k-window with decaying weights. This non-recursive design allows for future integration of biologically inspired constraints, such as limiting transitions based on sleep stage dynamics. The WES is calculated as:

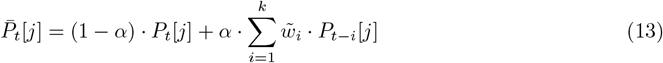

where:

- *α* ∈ [0, 1] is the smoothing parameter,
- *k* is the smoothing window size,
- *ŝ*_*t*−*i*_ = arg max_*j*_ *P*_*t*−*i*_[*j*] is the most probable class at time *t* − *i*,
- *P*_*t*−*i*_[*j*] is the predicted probability of class *j* at *t* − *i*,
- 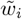 are the normalized exponential decay weights:

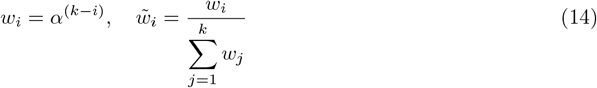

As mentioned, we also briefly investigate applying domain knowledge of physiologically plausible sleep transition constraints into the weighted exponential smoothing (WES) framework during evidence accumulation by modulating the contribution of each past prediction with a normalized transition prior matrix. The Transition-Constrained Weighted Exponential Smoothing (TC-WES) posterior is computed as:

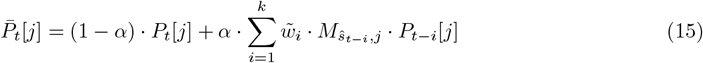

where: *M*_*i,j*_ ∈ [0, 1] is the normalized transition prior matrix, estimated empirically,

To compute *M*_*i,j*_, we avoided using transition frequency matrices, as they forced the model to converge toward the population’s average sleep pattern, ignoring individual differences. We tested this approach early on but found it overly biased and restrictive.

Instead, we built individual binary transition matrices for each of the 29 healthy subjects, where transitions with frequency *<* 1 were marked as disallowed (0) and others as allowed (1). These were averaged and normalized to create a continuous, subject-agnostic prior that reflects how *possible* a transition is, rather than how *often* it occurs; preserving biological plausibility without over-constraining the model.

While this work focuses on the unconstrained WES formulation, we also evaluated TC-WES. TC-WES yielded a noticeable improvement for one subject (*>*5% increase in performance), suggesting potential benefits in certain cases. However, the overall performance difference across subjects was marginal (others exhibited *<*1% change), indicating that hard constraints may also over-regularize or prevent recovery from early misclassifications, particularly when the model output itself is noisy or inaccurate.

#### Moving Average (MA)

This method averages the predictions over a sliding window of the past *k* timesteps:

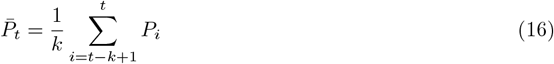

Unlike symmetric smoothing techniques that consider both past and future values, we adopt a causal formulation suitable for real-time applications by averaging only the previous *k* epochs. All past values within the window are weighted equally (i.e., unweighted averaging).

#### Kalman Filter (KF)

We applied a simplified Kalman filter to smooth the model’s posterior probabilities. The smoothed estimate 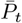 is recursively computed using a constant Kalman gain factor *K*_*t*_, determined by system and observation noise parameters *Q* and *R*:

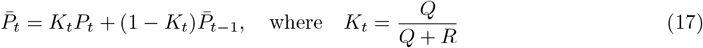

This formulation can be interpreted as an adaptive exponential smoothing mechanism, where *K*_*t*_ governs the relative influence of the current posterior *P*_*t*_ versus the past estimate 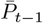. Although we do not explicitly model state dynamics or incorporate external observations, this recursive update structure captures temporal continuity and provides a lightweight, real-time smoothing strategy for closed-loop applications.

For smoothing methods, we have evaluated classification performance, temporal entropy and stability, delay, mismatch ratio, computational time, and trigger reliability, and demonstrate the practicality and robustness of each method in subject-independent, real-time settings.

All methods were applied to the softmax output sequence to generate temporally smoothed posterior probabilities 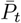, from which the final class predictions *ŷ*_*t*_ were derived.

Each smoothing method introduced one or two hyperparameters (e.g., *α* for ES, *α* and window size *k* for WES, window length *k* for MA, noise parameter *Q, R* for Kalman). We performed a 63-point individual grid search over plausible ranges of these parameters using the 5-subject validation set. The criteria for selection were maximizing the macro F1-score across all sleep stages on the validation set, as well as minimizing the mean temporal entropy and average delay (defined below) to encourage output stability. The optimal parameters (See Table. 2) provided a good balance of smoothing vs. responsiveness and were fixed for evaluating on the test set.

### 2.4. Evaluation Metrics

We assessed model performance by evaluating both classification accuracy and temporal stability using two complementary categories of metrics: *performance metrics* and *stability metrics*. Figure 2A, B shows the concept illustration of stability metrics calculation.

**Figure 2.**
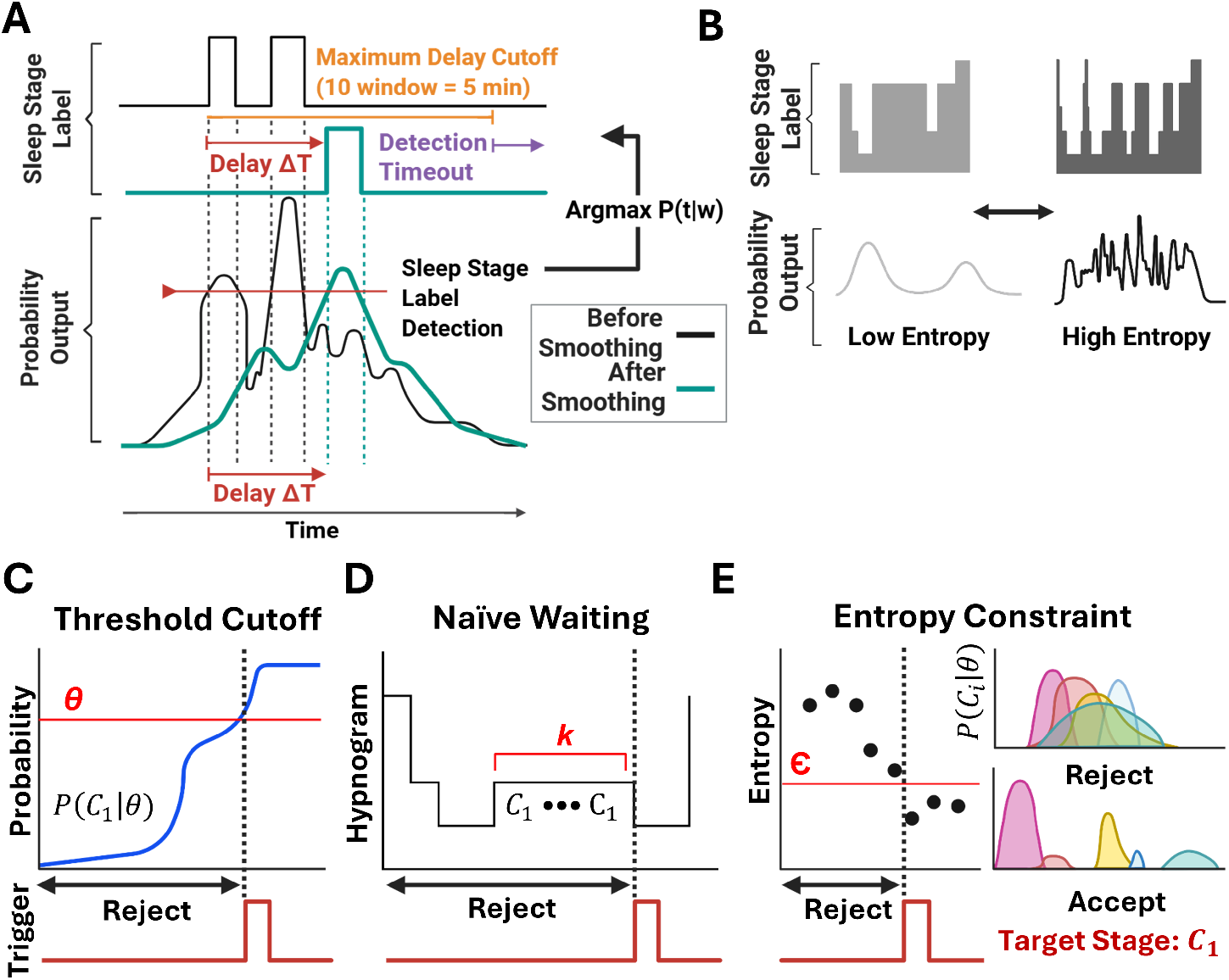
Concept Illustration of Stability Metrics and Various Conservative Logic. **(A, B)** Stability Evaluation Metrics Concept Illustration **(A)** Delay metric concept illustration. Raw (black) and smoothed (teal) posterior probabilities around a true sleep-stage transition (vertical dashed line) are shown for one class. An ANN’s posterior probability output is used to represent the standard decoder; after smoothing, sleep labels are detected based on the smoothed probabilities. Typically, Sleep labels are assigned by taking the *arg max* over five-stage probabilities at each time point. The detection delay ΔT is measured as the interval between the physiologist-annotated transition (red arrow) and the model’s smoothed-label detection (cyan arrow). If ΔT exceeds the maximum cutoff (10 windows = 5 min), a detection timeout (purple arrow) is declared, indicating a mismatch to meet real-time requirements in a closed-loop BCI system. This mismatch includes both model classification failure and detection delay timeout. The detailed definition and calculation of the mismatch ratio and delay can be found in Section 2.4.2. Averaging ΔT across all transitions quantifies each smoothing method’s impact on responsiveness versus conservatism. **(B)** Stability concept metric via temporal entropy. To capture label stability, we compute Shannon entropy of the five-class posterior distribution within non-overlapping 10-epoch (5 min) windows and average these values to yield a temporal-entropy score. Low entropy (left) corresponds to concentrated, unambiguous posteriors, ideal for stable BCI feedback and monitoring, whereas high entropy (right) reflects oscillatory, entangled probabilities that lead to unstable label assignments and might increase false positives. **(C, D, E)** Conceptual diagram of various conservative control logics compared to enhance trigger precision for targeted-stage stimulation following posterior smoothing. **(C)** *Threshold Cutoff:* A trigger is activated only when the target stage probability *P* (*C*_1_ | *θ*) exceeds a high threshold *θ*, ensuring conservative activation. **(D)** *Naive Waiting:* The system waits until the target stage *C*_1_ is maintained consistently for *k* consecutive windows before triggering, relying on temporal consistency. **(E)** *Entropy Constraint:* Our novel method rejects triggers when the Shannon entropy of the posterior exceeds a complexity threshold *ε*, indicating unstable or ambiguous stage predictions. The red square wave represents the final accepted trigger points under each logic.

#### 2.4.1. Model Performance Metrics

##### Precision, Recall, and F1-Score

We computed precision, recall, and F1-score for each individual sleep stage class. When to provide an overall assessment robust to class imbalance (e.g., the underrepresentation of N1), we report the *macro-averaged* F1-score, calculated as the unweighted mean of F1 across all classes. The F1-score is defined as the harmonic mean of precision and recall:

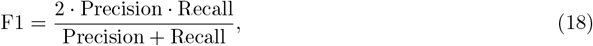

Where 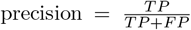, and 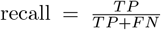. Where TP indicated true positives, and FP indicated false positives. Improvements in F1 indicate more accurate classification with fewer missed or false detections, especially useful in sleep staging with naturally imbalanced stage distributions.

##### Overall Accuracy

We additionally report overall accuracy as a baseline reference metric, though its interpretability is limited in imbalanced settings.

#### 2.4.2. Model Stability Metrics

##### Temporal Entropy

To quantify temporal consistency in model predictions, we introduce *temporal entropy*, defined as the average Shannon entropy of the predicted posterior distributions across all timepoints. For each epoch *t*, the entropy is computed as:

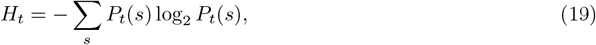

where *P*_*t*_(*s*) is the model’s probability output for stage *s* at time *t*. A lower average entropy indicates sharper and more confident predictions, whereas higher entropy suggests instability or indecision (e.g., an output like [0.4, 0.3, 0.3] has higher entropy than [0.9, 0.1, 0.0]). We report the mean entropy across all epochs.

##### Average Detection Delay

To evaluate responsiveness to stage transitions, we compute the average delay in correctly identifying the onset of sleep stages. Let *t*_true_ be the true start epoch of a target stage transition (e.g., REM→ Awake), and *t*_pred_ be the model’s first confident prediction of the same stage transition. Then the delay is defined as:

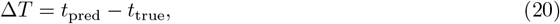

measured in a window unit and averaged across all occurrences of the target stage transitions. A smaller delay indicates faster reaction to stage onsets, which is especially important in real-time neuromodulation systems.

##### Latency-Bounded Mismatch Ratio

The *Latency-Bounded Mismatch Ratio* quantifies how often the model fails to detect true sleep stage transitions within a predefined temporal window (5 minutes = 10 windows). 5 min is selected due to its common usage for real-time monitoring. Formally, for each annotated transition, we consider it a mismatch if the model does not produce a correct prediction within the allowed maximum delay. This metric complements delay by capturing timing *failures*, not just timing *errors*. It is computed as:

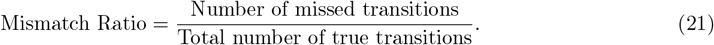

While a low average delay may suggest timely responses, a high mismatch ratio indicates that those responses often fall outside the acceptable time frame (bounded latency), making the detection unreliable for real-time applications. Taken together, delay and mismatch ratio provide a more complete picture of temporal fidelity than recall alone.

#### 2.4.3. Noise Robustness Testing

To evaluate the model’s robustness to noise, we added Gaussian noise with varying standard deviation from 0 to 1 to the input signals. The Gaussian noise used in the test is defined as:

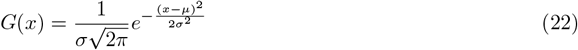

Here, *µ* is the mean and *σ* is the standard deviation of the Gaussian distribution. We applied this noise across a range of *σ* values to systematically test how the models behave under increasingly noisy conditions.

#### 2.4.4. Confusion Metrics Comparison

To quantitatively compare the performance of the smoothed model and the raw (non-smoothed) model under noise, we computed the difference in their confusion matrices for each test subject. The mean and standard deviation of these differences were used to analyze the consistency of the model behavior across subjects.

With test subjects (*N* = 4), the Delta Confusion Matrix is calculated as:

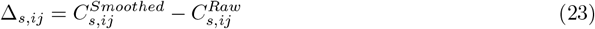

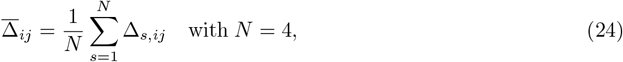

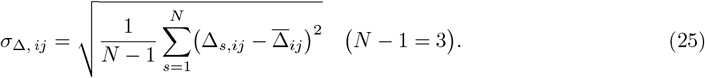

These metrics allowed us to capture not only the average performance difference between the two models but also how consistently those differences appeared across individual test subjects.

### 2.5. Conservative Triggering Control Logic

Having demonstrated overall improvements in model performance and output stability, a natural next question arises: can we further maximize precision, even at the cost of recall? In real-time closed-loop stimulation systems, minimizing false positives is paramount [7]. Which makes precision the most critical performance metric and widely studied to maximize [8,54]. While smoothing improves temporal stability, it does not explicitly guarantee the highest precision [55, 56].

Therefore, our second objective was to evaluate conservative decision rules (control logic) as a complementary safeguard against false or unstable outputs in real-world systems, to maximize stimulation precision for medical safety, and to assess whether smoothing techniques could enhance the reliability of triggering. In sleep medicine practice and research, it is common to impose ad-hoc criteria to ensure that a sleep stage is “truly” reached before responding. For example, conventional protocols often mandate a waiting period of 5–10 minutes after detecting a target stage before stimulation. One study implemented a 10-minute delay following continuous NREM detection to ensure stage stability [34], while another recommended a 15-minute delay after N3 onset for optimal efficacy [35]. Although such naïve waiting minimizes false positives, it also introduces substantial delays that may cause the system to miss brief yet valid stimulation windows—particularly problematic for transient stages like N1 or phasic REM [57].

Other strategies, such as probability thresholding [58, 59], activate stimulation only when the classifier’s confidence in a target stage exceeds a high cutoff (e.g., *P* (N2) *>* 0.7), treating lower-confidence predictions as noise and ignoring them [31–33]. While this can improve precision, it risks reduced sensitivity and vulnerability to output fluctuations. More recently, entropy-based measures have been explored as indicators of staging confidence: for instance, multi-scale entropy (MSE) analysis has shown that EEG signal entropy monotonically decreases from wake to deep sleep [60], suggesting that high output entropy may reflect ambiguity in classification and signal instability [61]. However, these insights have yet to be operationalized into simple, real-time control policies.

To address this gap, we systematically evaluated three representative control logics following posterior smoothing: (1) *Threshold Cutoff*, (2) *Naïve Waiting*, and (3) *Entropy Constraint*

We further introduce a hybrid control logic, the logic combination of the two, *threshold AND entropy constraints*, hypothesizing that this conjunction filters spurious detections more effectively than either criterion alone. This hybrid logic turns out to have the highest trigger precision by requiring both high confidence and low ambiguity. The algorithm is shown below. The conceptual illustrations in Fig. 2C–E summarize each logic’s conservative nature and operational impact on trigger decision timing.

To ensure generalizability across unseen subjects, all control logic parameters were fixed and not adaptively tuned. The threshold cutoff triggered stimulation only when the posterior probability of the target stage exceeded a clinically reliable range (*θ* = 0.7–0.8) [31–33]. The naïve waiting logic enforced temporal consistency by requiring stage persistence over three consecutive windows (1.5 minutes), a compromise between medical standards (typically ≥ 10 minutes) [34, 35] and the short durations of transient stages like N1 or phasic REM [36]. The entropy-based constraint suppressed triggers when the Shannon entropy of the posterior exceeded *ε* = 1.0, indicating high uncertainty or ambiguity in classification. All three logics were tested as non-adaptive, subject-independent policies designed to trade recall for higher precision in real-time settings.

#### Hybrid Control Trigger Algorithm

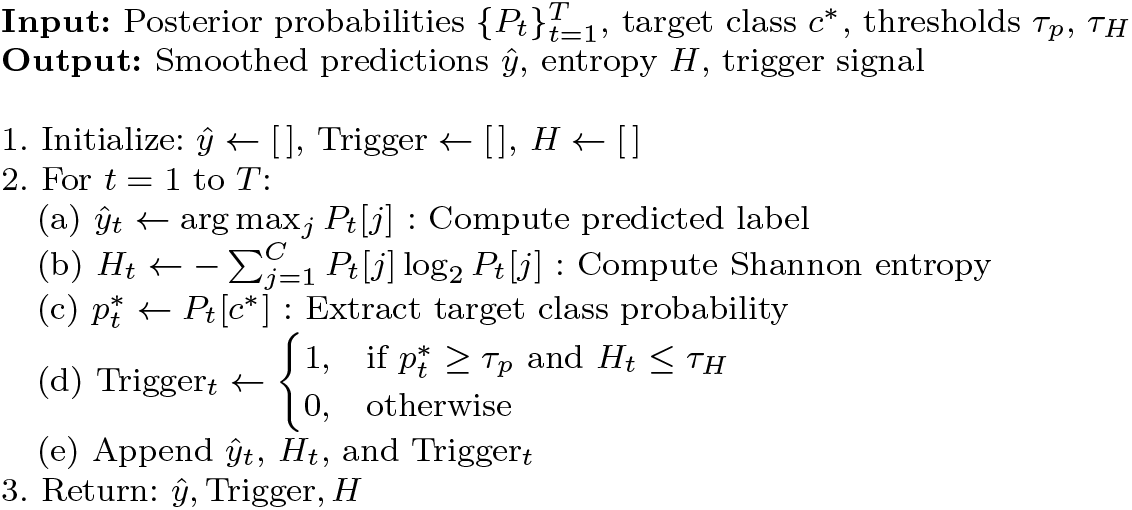

## 3. Results

The proposed post-processing pipeline consists of two key components: (1) temporal smoothing, aimed at enhancing classification performance by stabilizing the output probabilities; and (2) control logic, designed to maximize the precision of stimulation triggers of any targeted stage during sleep by enforcing stricter stage-confirmation criteria.

We hypothesized that temporal smoothing of model outputs would enhance prediction stability and overall classification performance by reducing spurious stage transitions, though potentially at the cost of responsiveness. Improving both interpretability and reliability in closed-loop systems is crucial where subjects are unconscious and predictions are inherently noisy. While smoothing may reduce responsiveness, it is likely to improve signal consistency by filtering out oscillatory noise that would otherwise lead to false positives, wasted energy, and make it harder to monitor and interpret. To test this, we evaluated four smoothing methods for their impact on predefined stability and performance metrics, identifying optimal smoothing intensities that generalize across individuals, which is a novel approach with implications for real-time system tuning. We additionally validate the robustness of temporal smoothing by testing on diverse synthetic noise conditions.

We further hypothesized that applying smoothing before control logic would amplify the precision gains of those strategies. Indeed, we found that post-smoothing control logics performed substantially better than on raw outputs. Through this stabilized setting, control triggers become more reliable due to reduced temporal noise. Among all strategies, a hybrid logic combining entropy and threshold constraints proved most effective: achieving over 96% precision for N2/N3 and up to 86% for REM triggers, while maintaining recall in the 75–88% range. This represents a substantial improvement over the raw model, which exhibited frequent false positives, especially during REM (precision ∼ 66%).

In addition, we introduce a transition presence matrix to capture individual-specific stage dynamics and investigate its potential as a domain-specific gain by applying it during the evidence accumulation process. This method enables potential applications in personalized or pathology-aware post-processing (e.g., for insomnia patients with atypical transitions). Importantly, the entire framework, including smoothing and control logic, is computationally lightweight, with runtime under 0.1 seconds per epoch on a standard laptop, making it feasible for deployment in real-time wearable or bedside systems.

Together, these findings define a robust, interpretable, and classifier-independent post-processing pipeline that significantly improves the reliability and efficacy of real-time sleep staging in closed-loop stimulation control.

### 3.1. Performance Trade-Offs for Different Smoothing Methods

To assess whether smoothing can enhance the stability and reliability of real-time sleep stage predictions, we first examined how four different smoothing methods affect both model performance and temporal stability. While smoothing can suppress noisy posterior fluctuations, excessive smoothing may hinder responsiveness to true state transitions. Thus, in this subsection, we aim to answer two questions: Is there an existence of an optimal smoothing range that maximizes performance while maintaining acceptable latency? - and if it is generalizable and consistent, in what range? Our baseline neural network model achieved an overall accuracy of 78.2% across five sleep stages on a 29-subject dataset, with class-specific F1-scores of 84% (Wake), 38% (N1), 83% (N2), 85% (N3), and 80% (REM) by using a 29-group K-fold. This base model was used as the baseline for all later analyses. Our focus was not on improving the deep learning model’s architecture, but on testing post-processing methods that work independently of the model itself. Importantly, parameter tuning and evaluation metrics for each method were exclusively performed using the validation dataset (N=5), while the test set (N=4) was reserved for one-shot real-time evaluation without any adaptation or optimization.

We evaluated four smoothing methods: Moving Average (MA) [51], Exponential Smoothing (ES) [22, 50], Kalman Filtering (KF) [52, 53], and our proposed Weighted Exponential Smoothing (WES). Each method was applied across a range of normalized smoothing intensities (0 = no smoothing to 1 = maximum smoothing). We then quantified their effects on F1-score for each sleep class, average delay, entropy, and mismatch ratio.

As shown in Fig. 3, WES and ES outperformed the other two methods, maintaining steady and high F1-scores across intensities with minimal degradation at higher smoothing. WES (A) maintains the most consistent performance across all intensities; ES (B) performs comparably until a sharp drop beyond 0.8; MA (C) shows generally lower F1-scores and decreases from the beginning; and Kalman filtering (D) exhibits increasing fluctuation (unstable prediction) at high smoothing intensities after 0.5; For all methods, The N1 stage (red) remains the most challenging to classify, consistent with previous studies [27, 62], reflecting its low baseline separability of the base decoder itself. To complement F1-score analysis, we assessed how stability metrics – Entropy, Delay, and Latency-Bounded Mismatch Ratio(REF) varied with smoothing intensity. In Figure 4A–C, Entropy, reflecting output uncertainty and complexity in the model’s posterior distribution, decreased monotonically across all methods as smoothing increased. WES and ES exhibit a gradual entropy decline, suggesting they avoid over-simplification while still improving stability. MA yields the lowest final entropy with the lowest F1-Score (See Fig. 3). This indicates that while MA produces highly stable outputs, it may suppress too much temporal detail. Meanwhile, the mismatch rate, used as a proxy for missed or overly delayed transitions, increased with stronger smoothing (Fig. 4B), with MA again showing the worst latency. In contrast, WES maintained lower timeout and delay even at higher intensities (Fig. 4C). These trends underscore the trade-off between prediction stability and responsiveness and support using a physiologically constrained sweet spot (entropy *<* 0.7, delay *<* 5 epochs) to guide optimal parameter selection.

**Figure 3.**
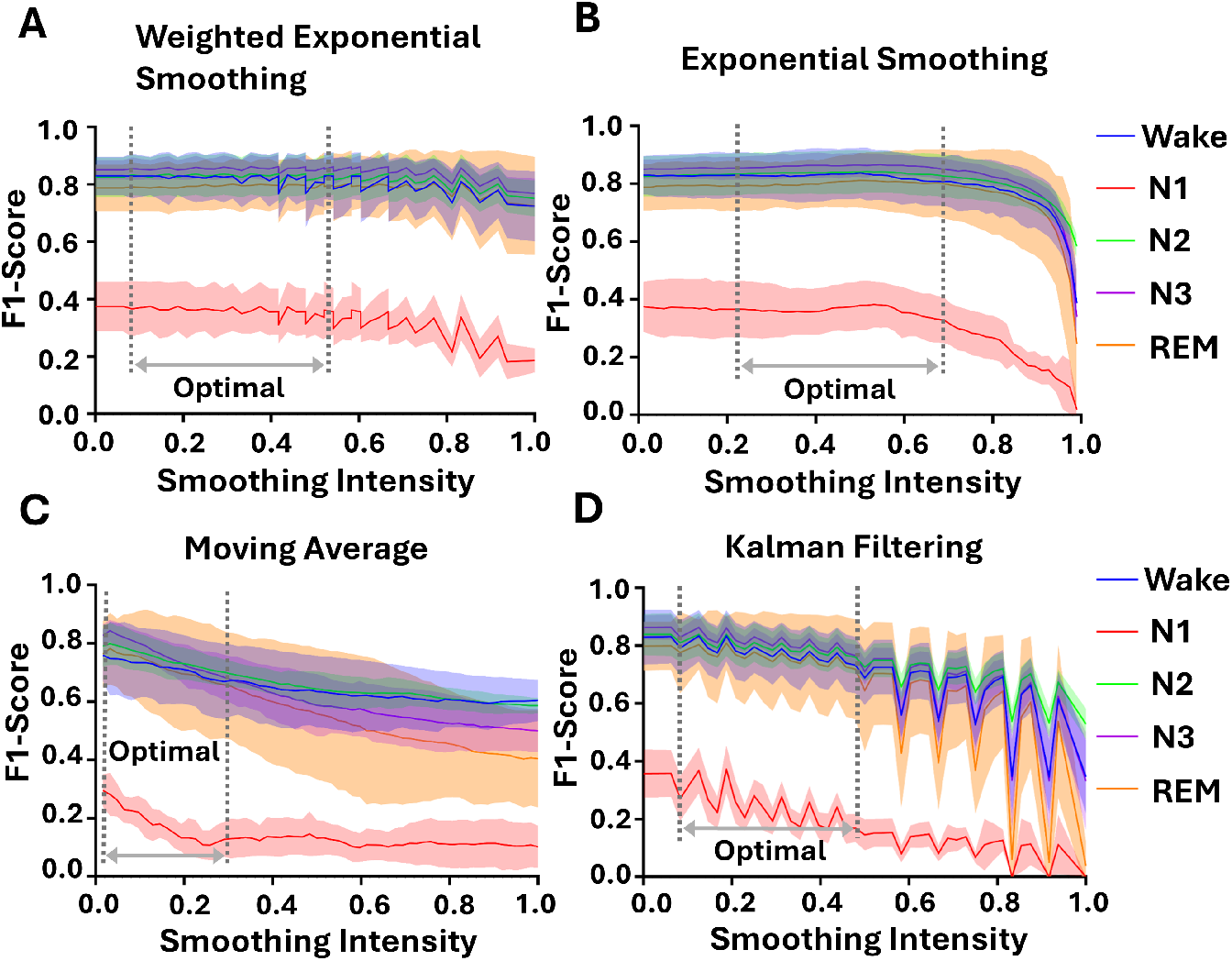
F1-score Versus Normalized Smoothing Intensity for Five Sleep Stages (Wake: blue; N1: red; N2: green; N3: purple; REM: orange) Using Four Different Smoothing Techniques. **(A)** Weighted Exponential Smoothing (WES), **(B)** Exponential Smoothing (ES), **(C)** Moving Average (MA), and **(D)** Kalman Filtering (KF). The x-axis spans from 0 (raw posterior outputs) to 1 (maximum smoothing). Shaded “optimal” regions indicate the range of smoothing intensities that yield both high and stable F1-scores, corresponding to minimal prediction entropy (stability), delay, and mismatch rate-thus representing the optimal trade-off. F1-score, which balances precision and recall in class-imbalanced data, is averaged across the validation sleep dataset from five subjects.

**Figure 4.**
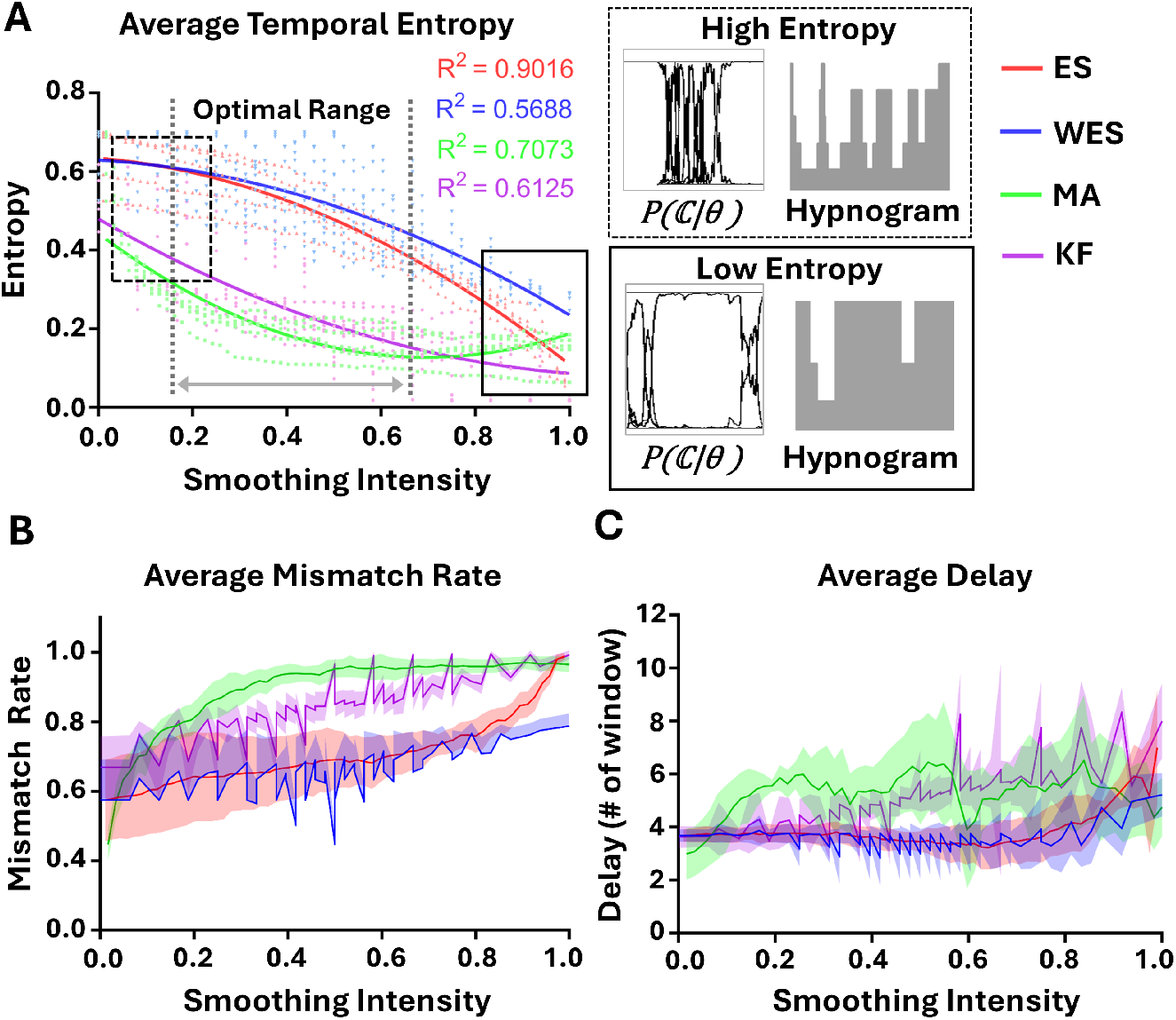
Performance of Stability Metrics (Entropy, Mismatch Rate, and Delay) Across Normalized Smoothing Intensity for Four Smoothing Techniques. **(A)** Average Temporal Entropy vs. Normalized Smoothing Intensity. Temporal entropy was computed from the five-stage posterior distribution in each 10-epoch window (5 min), consistent with maximum delay cutoff (See Fig. 2A), is averaged across all sleep stages and five validation subjects, then plotted against normalized smoothing intensity (0 = raw, 1 = max). Second-order polynomial fits yield *R*^2^ of 0.9016 for Exponential Smoothing (red), 0.5688 for WES (blue), 0.7073 for Moving Average (green), and 0.6125 for Kalman Filtering (purple). Entropy falls monotonically with stronger smoothing, reflecting progressively simpler hypnograms (right panels) and more stable posterior profiles. WES shows the most gradual entropy drop, while MA attains the lowest overall entropy (most stable, monotonic outputs). **(B)** Average mismatch rate across all sleep stages vs. Normalized smoothing intensity, defined as proportion of corresponding transitions (See Method section 2.4.2 for more detail) within bounded latency Δ*T <* 10 windows (5 min). MA exhibited the highest mismatch rate, while WES showed the lowest. ES remained comparable to WES until approximately 0.8, after which its mismatch rate increased sharply. **(C)** Average detection delay (Δ*T* ) across all sleep stages vs. Normalized smoothing intensity. Delay rises gradually with smoothing strength; WES and ES maintain shorter delays than MA and KF.

Together, these results identify an “optimal range” of smoothing intensity that simultaneously enhances F1-score and reduces complexity, while keeping delay within acceptable limits.

### 3.2. Optimal Smoothing Intensities and Validation Performance

Having observed an optimal trade-off region between stability and performance, we next sought to identify parameter configurations that maximized macro F1-score under biologically grounded constraints. Specifically, we selected smoothing intensities that (1) reduced entropy below 0.7 (since most of performance falls under entropy below 0.7) and (2) maintained detection delay under 5 epochs (2.5 minutes). These criteria ensure clinical responsiveness while avoiding false alarms in real-time systems. These optimal values and normalized intensity of the given parameter set are summarized in Table 2 (the optimal parameter table), confirming that a moderate smoothing level is best for all methods. We report that an optimal amount of smoothing dramatically reduces the entropy (or uncertainty) of the predicted hypnogram (decreased up to 29%), yielding cleaner state transitions and improving overall accuracy by up to 2% in a model-agnostic manner. In deep learning, especially on high-performing models (e.g., 85–90% baseline), even a 1–2% absolute increase is non-trivial, often requiring architectural changes, hyperparameter tuning, additional data, or extended training; making our lightweight, generalizable gain particularly noteworthy. This 2% translates to nearly 10 minutes (20 windows) of reduced false positives for 8 hours of sleep per night.

**Table 2.** Optimal Parameters of Each Smoothing Method Derived from Validation Set (N=5)^*†*^.

| Smoothing Method | Optimal Parameters<br>*Mean ± SD* | Smoothing Intensity<sup>†</sup> |  |
| --- | --- | --- | --- |
|  |  | Mean | SD |
| WES | $\alpha = 0.467 \pm 0.073$, $k = 3.80 \pm 0.837$ | 0.408 | 0.079 |
| ES | $\alpha = 0.393 \pm 0.198$ | 0.393 | 0.198 |
| MA | $k = 3.20 \pm 1.789$ | 0.052 | 0.029 |
| KF | $Q = 0.006 \pm 0.001$, $R = 0.01 \pm 0.000$ | 0.225 | 0.071 |

<sup>†</sup>Smoothing intensity values were normalized to $[0, 1]$ for comparison across methods: WES was normalized by averaging the scaled smoothing factor ( $\alpha \in [0.01, 0.9]$ ) and window size ( $k \in [1, 9]$ ). ES intensity corresponds to the normalized $\alpha$ in the range $[0.01, 0.99]$ . MA (Moving Average) was scaled based on window size $k \in [1, 63]$ . For Kalman Filtering, intensity was computed as the average of normalized $Q \in [10^{-5}, 10^{-2}]$ and $R \in [10^{-2}, 1.0]$ , where smaller $Q$ and larger $R$ imply stronger smoothing, respectively.

For each smoothing method, we computed the optimal parameter set by averaging across the five validation subjects. WES, our proposed method, exhibited consistent optimal parameters: *α* = 0.467 ± 0.073, *k* = 3.80 ± 0.837. ES also converged near *α* = 0.393 ± 0.198, while MA showed more variability in *k* = 3.20 ± 1.789, and KF had tightly clustered values for *Q* = 0.006 ± 0.001 and *R* = 0.01 ± 0.000. Notably, the normalized smoothing intensities associated with optimal performance were remarkably similar across subjects, particularly for WES and ES (mean ≈ 0.4 ∼ 0.5), suggesting generalizability across individual variations.

To assess the benefit of optimal smoothing, we compared classification performance metrics before (raw output) and after applying these optimal smoothing parameters (see Fig.5). Across the validation set, all performance metrics: f1-score, precision, and recall, showed improvements after smoothing both in the subject level (see Table. 3, and the group level. To quantify the significance of these improvements, we conducted paired t-tests (N = 5 subjects), and observed gains were statistically meaningful.

**Figure 5.**
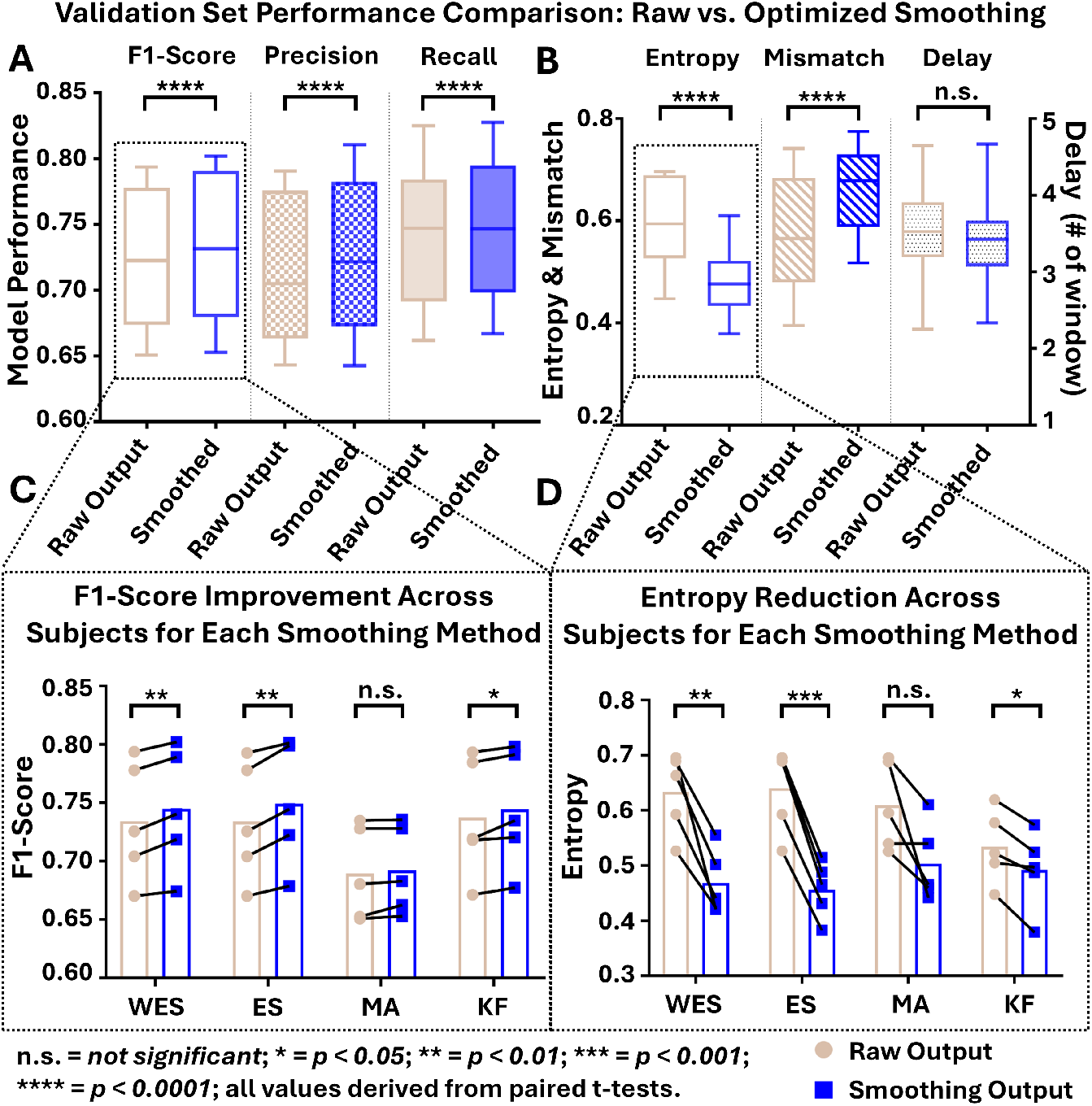
Validation Set Comparison of Model Performance (f1-score, precision, recall) and Stability Metrics (entropy, mismatch ratio, delay) Before (raw output) and After Smoothing Across Four Smoothing Methods (N = 5). **(A)** Box plots of macro-averaged f1-score, precision, and recall across five sleep stages. After applying optimal smoothing, all methods showed consistent improvements. Although these increases were modest due to strong baseline performance and limited sample size, the trend was consistent across all smoothing methods. **(B)** Comparison of stability metrics: entropy, mismatch ratio, and delay. All methods significantly reduced entropy, both at the group and subject levels, indicating greater output consistency. Mismatch ratio slightly increased post-smoothing (e.g., WES: +7.9%), a trade-off reflecting suppression of brief predictions. Delay either decreased or remained stable (e.g., WES: 0.25 window). Details can be found in the supplementary information. **(C)** Per-subject comparison of F1-score before and after smoothing shows that all five subjects experienced improvement across all methods, demonstrating robustness of the smoothing effect. Each connected line of raw output to smoothing output indicates the same subject. WES improved performance from 0.734 ± 0.051 to 0.745 ± 0.052 (+1.5%), ES from 0.734 ± 0.051 to 0.74 ± 0.052 (+2.0%), MA from 0.689 ± 0.040 to 0.692 ± 0.038 (+0.4%), and KF from 0.737 ± 0.051 to 0.744 ± 0.051 (+0.9%). Paired t-tests revealed significant improvements in WES and ES (*p* = 0.0061, *p* = 0.0051, respectively), moderate significance for KF (*p* = 0.0346), and no statistical significance for MA (*p* = 0.1669), although most subjects showed improvement. Statistical significance across sessions is marked (*ns*: not significant; ^∗^*p <* 0.05; ^∗∗^*p <* 0.01; ^∗∗∗^*p <* 0.001, ^∗∗∗∗^*p <* 0.0001). When aggregating all four smoothing methods and comparing them to raw predictions, we observed a highly significant group-level effect (*p <* 0.0001). This trend was consistent across macro F1-score, macro precision, and macro recall for all sleep stages, further confirming the robustness of the smoothing effect. Combined analysis across all methods again demonstrated a highly significant group-level difference (*p <* 0.0001). **(D)** Corresponding entropy reduction for each subject confirms the consistent stabilization effect, with WES and ES showing the most pronounced drops. WES reduced entropy from 0.634 ± 0.072 to 0.468 ± 0.059 (26.2%), ES from 0.640 ± 0.077 to 0.456 ± 0.051 (28.8%), MA from 0.609 ± 0.081 to 0.503 ± 0.071 (17.4%), and KF from 0.534 ± 0.067 to 0.492 ± 0.071 (7.9%). WES and ES showed highly significant reductions (*p* = 0.0049, *p* = 0.0002), KF showed a modest yet significant drop (*p* = 0.0136), and MA showed no statistical significance (*p* = 0.0615). Combined analysis across all methods again demonstrated a highly significant group-level effect (*p <* 0.0001). *Note:* Smoothing parameters used are those previously optimized for each method (see Table 2).

Regarding macro F1-score, WES improved performance from 0.734 ± 0.051 to 0.745 ± 0.052 (+1.5%), ES from 0.734 ± 0.051 to 0.749 ± 0.052 (+2.0%), MA from 0.689 ± 0.040 to 0.692 ± 0.038 (+0.4%), and KF from 0.737 ± 0.051 to 0.744 ± 0.051 (+0.9%). Paired t-tests revealed significant improvements in WES and ES (*p* = 0.0061, *p* = 0.0051, respectively), moderate significance for KF (*p* = 0.0346), and no statistical significance for MA (*p* = 0.1669), although most subjects showed improvement. When aggregating all four smoothing methods and testing them against raw predictions, we observed a highly significant group-level effect (*p <* 0.0001).

Precision and recall followed a similar pattern and are reported in Table 3. For prediction entropy, smoothing yielded substantial reductions, reflecting enhanced temporal consistency. WES reduced entropy from 0.634 ± 0.072 to 0.468 ± 0.059 (26.2%), ES from 0.640 ± 0.077 to 0.456 ± 0.051 (28.8%), MA from 0.609 ± 0.081 to 0.503 ± 0.071 (17.4%), and KF from 0.534 ± 0.067 to 0.492 ± 0.071 (7.9%). WES and ES showed highly significant reductions (*p* = 0.0049, *p* = 0.0002), KF showed a modest yet significant drop (*p* = 0.0136), and MA showed no statistical significance (*p* = 0.0615). Combined analysis across all methods again demonstrated a highly significant group-level effect (*p <* 0.0001).

**Table 3.** Comparison of Validation Set Performance: Raw vs. Optimized Smoothing Method Across Sleep Stages (N=5)

of model performance metrics of each smoothing method (N = 5, Mean ± SD)*
|  | Macro F1-Score |  | Macro Precision |  | Macro Recall |  |
| --- | --- | --- | --- | --- | --- | --- |
|  | Raw | Smoothed | Raw | Smoothed | Raw | Smoothed |
| WES | 0.734±0.051 | **0.745±0.052** | 0.724±0.051 | **0.739±0.052** | 0.755±0.056 | **0.760±0.055** |
| ES | 0.734±0.051 | **0.749±0.052** | 0.724±0.051 | **0.744±0.054** | 0.755±0.055 | **0.764±0.054** |
| MA | 0.689±0.040 | **0.692±0.038** | 0.680±0.039 | **0.685±0.039** | 0.707±0.042 | **0.709±0.039** |
| KF | 0.737±0.051 | **0.744±0.051** | 0.730±0.053 | **0.737±0.051** | 0.756±0.054 | **0.762±0.053** |

*Result of model stability metrics of each smoothing method (N = 5, Mean ± SD)*
|  | Average Entropy |  | Mismatch Ratio |  | Average Delay |  |
| --- | --- | --- | --- | --- | --- | --- |
|  | Raw | Smoothed | Raw | Smoothed | Raw | Smoothed |
| WES | 0.634±0.072 | **0.468±0.059** | **0.587±0.119** | 0.666±0.094 | 3.753±0.351 | **3.503±0.433** |
| ES | 0.640±0.077 | **0.456±0.051** | **0.577±0.117** | 0.683±0.071 | 3.694±0.245 | **3.429±0.546** |
| MA | 0.609±0.081 | **0.503±0.071** | **0.475±0.047** | 0.683±0.071 | **3.089±0.923** | 3.429±0.546 |
| KF | 0.534±0.067 | **0.492±0.072** | **0.669±0.089** | 0.696±0.066 | 3.634±0.411 | **3.516±0.476** |

*\*Note: Bolded values indicate the best (most favorable) performance for each metric, not necessarily the largest or smallest numerical value. Raw refers to model output without smoothing, while Smoothed refers to performance after applying the smoothing method. Model performance metrics (F1-Score, Precision, Recall) represent macro-averaged values across 5 sleep stages and are reported as mean ± SD over 5 validation subjects. For these metrics, higher is better. Real-time evaluation metrics: Delay, Entropy, and Mismatch Ratio, are also reported as mean ± SD. Lower values indicate better performance for all three: Entropy (probabilistic complexity for monitoring and output stability), Delay (average temporal latency), and Maximum Latency ( $< 5min$ ) Bounded Mismatch Ratio.*

Delay remained unaffected by smoothing. For instance, WES showed a minimal change from 3.753 ± 0.351 to 3.503 ± 0.433 (*p* = 0.1106), indicating that improved stability did not come at the cost of temporal responsiveness.

However, mismatch ratio slightly increased (e.g., WES from 0.587 ± 0.119 to 0.666 ± 0.094; *p <* 0.0001), suggesting a more conservative model behavior, potentially trading off some temporal recall for stability. Overall, WES and ES achieved the most consistent improvements across all metrics, supporting the hypothesis that temporal smoothing can enhance prediction performance in a model-independent and physiologically-informed manner without introducing substantial latency.

### 3.3. Generalization of Smoothing Improvement on Unseen Subject

Since the optimal parameters seem similar across subjects, can we use this value to unseen test subject to expect a performance increment? To validate whether smoothing gains could generalize to previously unseen individuals, we applied the average optimal WES parameters (*α* = 0.5, *k* = 4) to all test sets (N=4), without further tuning. Results showed consistent, albeit modest, improvements in macro F1-score, precision, and recall after smoothing in the group level (F1-score: 0.740 ± 0.074 → 0.743 ± 0.093 (+0.3%), precision: 0.745 ± 0.053 → 0.760 ± 0.069 (+1.5%), recall: 0.778 ± 0.051 → 0.785 ± 0.005 (+0.7%)). On the subject level, two of the four subjects exhibited clear improvements (f1-score increase from 0.79 to 0.83 (+4%) for subject 2, and from 0.81 to 0.83 (+2%) for subject 4, other two subjects drop by 2% (from 0.65 to 0.63) and 1% (from 0.71 to 0.70); the overall trend suggests that our method provides stable performance enhancements even under one-shot test conditions. While smoothing improved precision and recall for all individuals, statistical comparisons using a paired t-test (N = 4) did not reach significance for macro F1-score, precision, or recall (all *p >* 0.05; *p* = 0.7306, 0.2452, *and* 0.5707, respectively). Given the small sample size and within-subject consistency, we interpret these trends as indicative rather than conclusive, yet the consistency in optimal parameter values offers promise for broader applicability.

To further illustrate the smoothing effect at the individual level, we visualized an 8-hour sleep record from Subject 2 in the test set. Subject 2 is selected within the test data due to its longest sleep time and balanced sleep stages. As shown in Fig. 6, smoothed prediction’s entropy overlays on the true hypnogram (Fig. 6A) reveal that high-entropy epochs align with physiologically ambiguous transitions, particularly at stage boundaries. Before smoothing (Fig. 6B, top), posterior probabilities were highly noisy and complex, displaying sharp fluctuations even during stable sleep states. After applying WES (Fig. 6B, bottom), predictions became more consistent and cleaner, matching the expert-scored hypnogram. This visual evidence confirms that smoothing not only improves quantitative metrics but also enhances interpretability and trust in stable system outputs.

**Figure 6.**
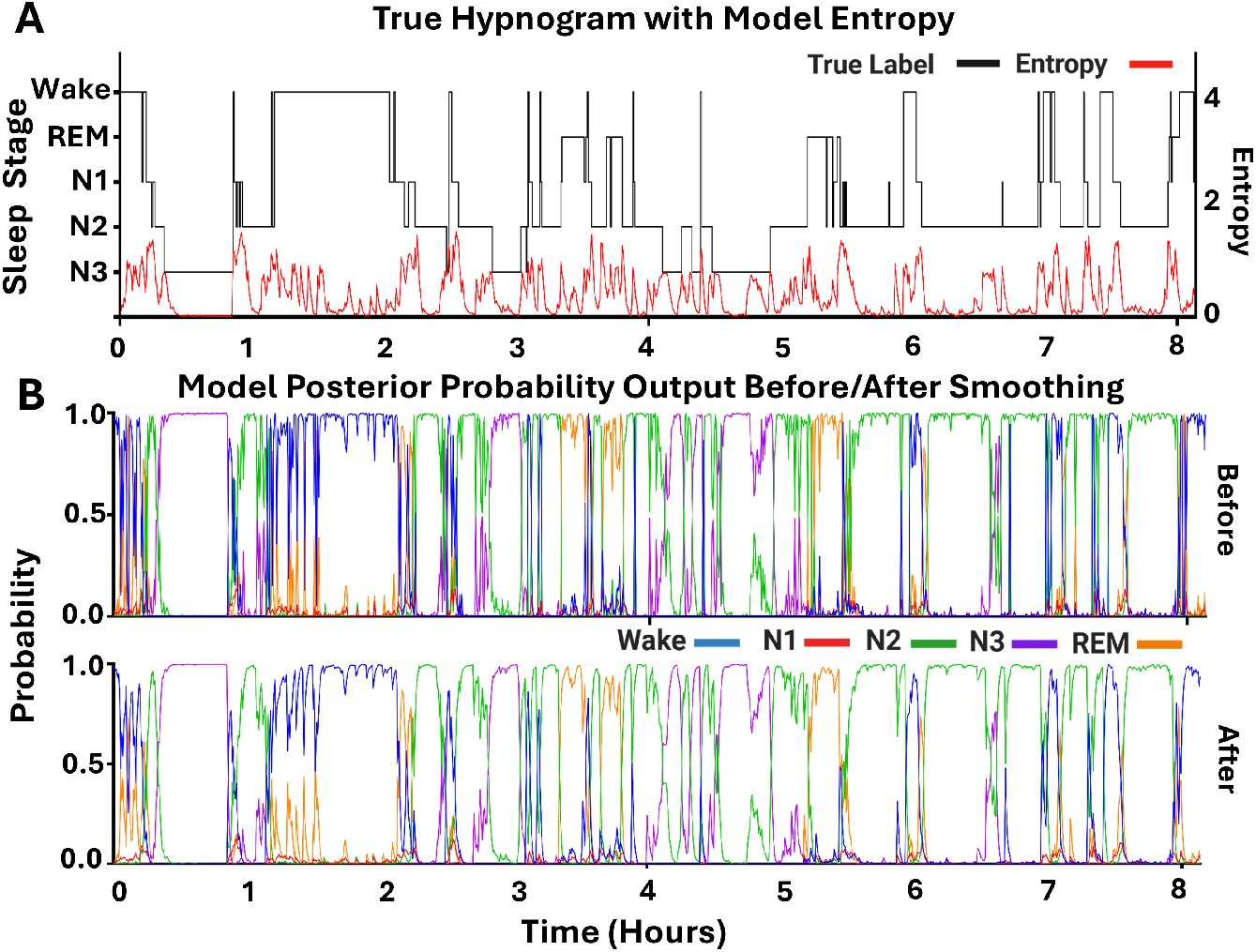
Impact of WES Smoothing on Model Entropy and Posterior Probabilities During Sleep Stage Classification on an Individual Test Subject. An eight-hour recording of subject 2 was selected to illustrate model behavior before and after smoothing. The representative subject was selected due to its longest sleep time and balanced sleep stages. **(A)** Expert-scored hypnogram (black) overlaid with the Shannon entropy calculated for each window of the smoothed posterior probabilities (red). Entropy peaks correspond to the epochs in which the classifier’s posterior distribution is most uniform—i.e., the model is less certain and finds it harder to discriminate among sleep stages. **(B)** Posterior probability outputs for each sleep stage, Wake (blue), N1 (red), N2 (green), N3 (purple), REM (orange), before smoothing (top) and after applying the WES smoothing method (bottom). Smoothing parameters (*σ* = 0.5, *window* = 4) were chosen based on the optimal performance in the validation set (See Table. 2). The WES smoothed probabilities showed reduced rapid and unstable fluctuations in the model posterior.

### 3.4. Robustness of Smoothed Outputs to Noise

We next investigate whether this smoothing strategy remains effective and robust under different noisy conditions that mimic real-world sensor imperfections or subject variability. To answer this, we performed a controlled experiment by injecting Gaussian noise (*σ* = 0 to 1.0) into the model’s posterior outputs and assessing classification performance before and after smoothing. A larger sigma indicates a higher noise scenario. Fig. 7A demonstrates that as noise increases, raw model predictions degrade rapidly in both precision and recall. In contrast, the WES smoothed outputs retain significantly higher performance consistently beyond *σ* = 0.2 to 0.3, with up to 10%-15% improvement in both metrics. Confusion matrix comparisons (Fig. 7 B-D) further show that smoothing corrects misclassifications across all stages except N1, which remains challenging. A delta mean heatmap analysis (See Equation 24) across four test subjects under Gaussian noise (*σ* = 0.5) reveals a notable increased occurrence in true positives stage detection: Wake (+13.50), N2 (+55.25), N3 (+24.50), and REM (+17.50). In contrast, most false positives were visibly reduced (highlighted in red), suggesting strong noise robustness. However, N1 showed a decrease in true positives ( −5.25), likely due to its transitional nature and the smoothing-induced suppression of brief stage fluctuations. Fig. 7 E-G illustrates that noisy raw hypnograms exhibit stage oscillations and false positives, while smoothed outputs preserve a clearer and more physiologically plausible sleep structure. Together, these results suggest that WES provides robustness against probabilistic instability induced by noise, further justifying its use in real-time closed-loop systems.

**Figure 7.**
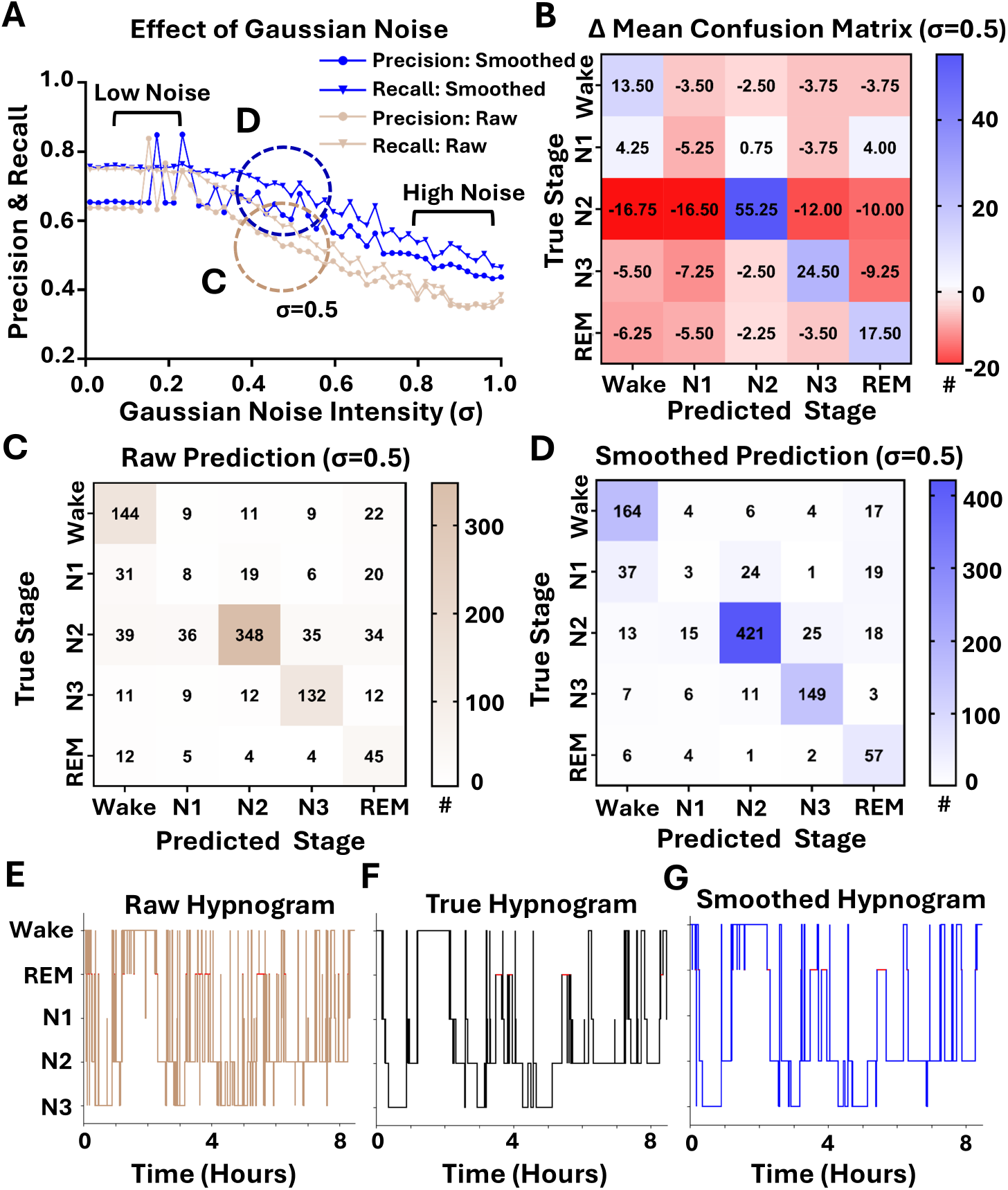
Impact of Gaussian Noise on Model Performance and Sleep Hypnograms Before and After WES Smoothing. Sleep recording from the test set was used for classification performance with added Gaussian noise. **(A)** Precision (circle) and recall (triangles) for raw outputs (beige) versus WES smoothed output (blue), plotted as a function of Gaussian noise intensity (*σ*) added to the model’s posterior probabilities. At low noise (*σ <* 0.3), raw and smoothed curves are similar; beyond *σ* = 0.3, smoothing yields a consistent 10%-15% improvement in both precision and recall (e.g., for *σ*=0.3, Accuracy: 0.775 → 0.855 (+18%), macro precision: 0.690 → 0.781 (+9.1%), macro recall: 0.734 → 0.812 (+7.8%)). # in color bar indicates the number of occurrences of sleep stage. **(B)** Mean of Difference (See Equation. 24) in the Confusion matrix across subjects at *σ* = 0.5. The difference heatmap in (B) was computed by subtracting the raw prediction heatmap (C) from the smoothed prediction output (D). Positive (blue) diagonal entries and negative (red) off-diagonals indicate that smoothing increases correct stage assignments and reduces missclassification for all sleep stages except N1. This result is consistent with all noise levels. **(C)** Confusion matrix for Subject 2 and *σ* = 0.5 using raw predictions **(D)** Corresponding confusion matrix after WES smoothing. **(E-G)** Hypnograms for Subject 2 at *σ* = 0.5: raw predicted staging (E), expert-scored true labels (F), and WES-smoothed staging (G). Raw predictions (E) exhibit rapid oscillations and numerous false positives; smoothing produces a more stable, true hypnogram-like sleep architecture (G).

### 3.5. Trigger Precision Enhancement via Control Logic Constraints

Building on the observed improvements in model performance and temporal stability, we next investigated whether simple control strategies could further enhance trigger precision. We hypothesized that applying our smoothing step prior to control logic methods would amplify their effectiveness, particularly by stabilizing predictions before constraint enforcement. To ensure robust deployment in real-time closed-loop systems, we explored simple, fixed control logics that prioritize trigger precision. By comparing threshold-cutoff, naïve waiting, and our proposed entropy-based constraints, we demonstrated that even under non-adaptive conditions, conservative control strategies can effectively reduce false positives.

As shown in Fig. 8, applying smoothing prior to control logic led to a clear and consistent increase in precision across all sleep stages, than directly applying control logic to raw output. Notably, the Hybrid Constraint (threshold + entropy) method achieved the highest precision in every case, while Naïve Waiting consistently underperformed. These findings support our hypothesis that smoothing helps stabilize model outputs, allowing subsequent control logic to more effectively suppress false positives. Furthermore, the improvement from Raw + Hybrid to Smoothed + Hybrid control logic: from 0.703 to 0.799 (Awake), 0.941 to 0.963 (N2), 0.968 to 0.981 (N3), and 0.724 to 0.861 (REM), demonstrates the strong benefit of combining probabilistic smoothing with conservative control logic —together as “post-processing” framework.

**Figure 8.**
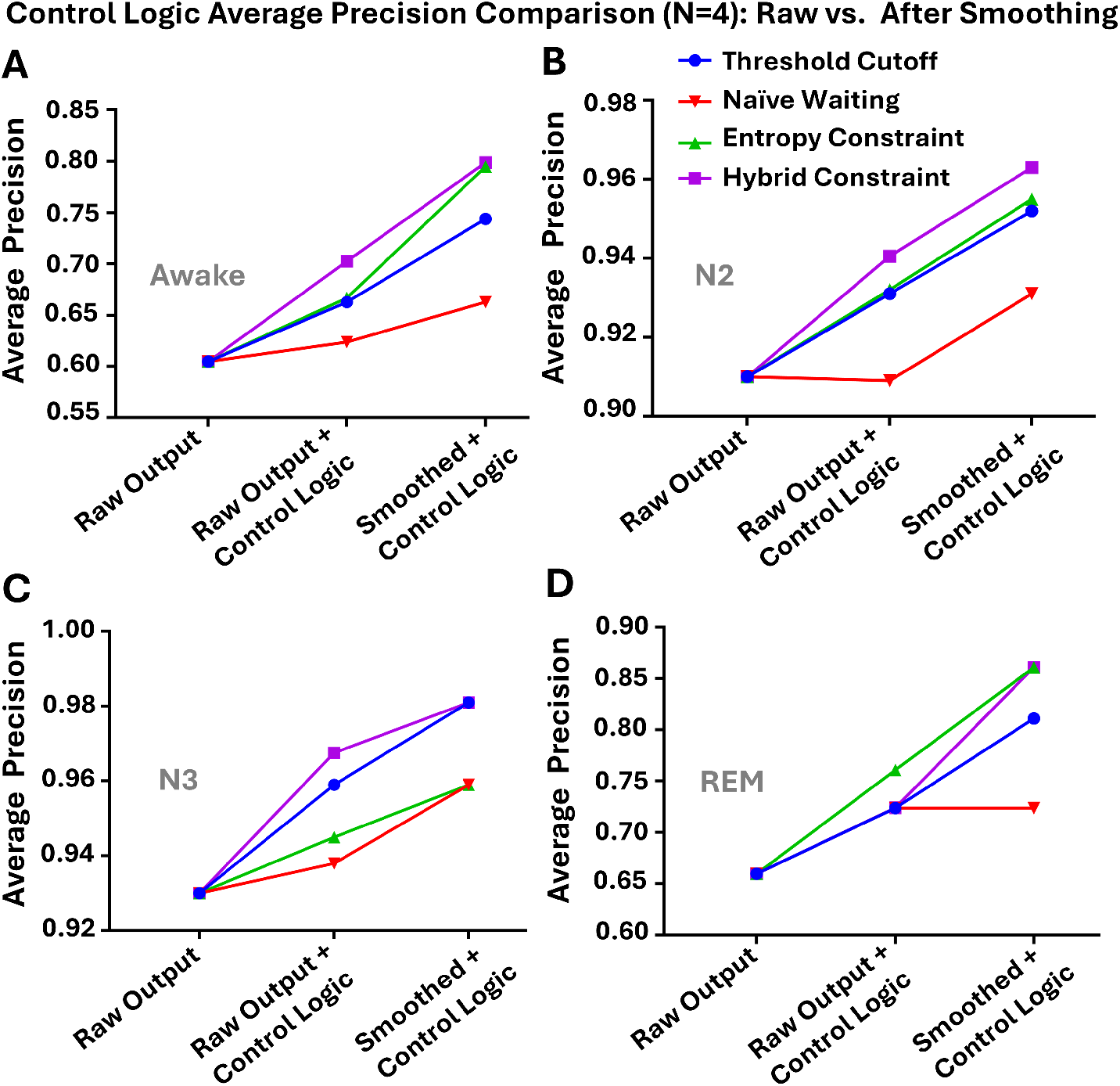
Control Logic Performance Across Sleep Stages (N=4). This figure illustrates how average precision changes depending on the target sleep stage (Awake, N2, N3, REM) when applying different control logic strategies to raw ANN model outputs. We compare three scenarios: the unmodified *Raw Output* (baseline model performance), applying control logic directly to the raw output (*Raw Output + Control Logic; Before*), and applying control logic after smoothing the model output (*Smoothed + Control Logic; After* ). Across all sleep stages, smoothing followed by control logic leads to improved precision, supporting the hypothesis that smoothing stabilizes the probability distribution, enabling control logic to more effectively suppress false positives. However, this may come at the cost of reduced recall (see Table. 4) Notably, the *Hybrid Constraint* strategy (purple) consistently achieves the highest precision across all stages, while the *Naïve Waiting* (red) method (a commonly used but overly simplistic approach) results in the lowest precision. The average precision scores (mean SD) from the unmodified ANN model are: Awake = 0.605 ± 0.256, N2 = 0.910 ± 0.041, N3 = 0.930 ± 0.043, REM = 0.660 ± 0.124. For *Raw + Hybrid Control Logic*, the precision scores are: Awake = 0.703 ± 0.201, N2 = 0.941 ± 0.029, N3 = 0.968 ± 0.027, REM = 0.724 ± 0.113. For *Smoothed + Hybrid Control Logic*, the precision scores further improve: Awake = 0.799 ± 0.164, N2 = 0.963 ± 0.015, N3 = 0.981 ± 0.019, REM = 0.861 ± 0.097. The above standard deviation(error bars) were omitted here for clarity.

Detailed information is summarized in Table 4, and smoothing consistently improved precision across all control logic methods. For example, in the Awake class, precision increased from 0.663 ± 0.220 (raw) to 0.744 ± 0.176 (+8.1%, smoothed). Similarly, N2 improved from 0.931 ± 0.034 to 0.952 ± 0.022 (+1.9%), N3 from 0.959 ± 0.022 to 0.981 ± 0.019 (+2.2%), and REM from 0.724 ± 0.107 to 0.811 ± 0.113 (+8.7%). These trends were consistent across all three constraint types (threshold, naïve waiting, and entropy constraint). Statistical comparisons using a paired t-test (N = 4) did not reach significance for most stages due to the small sample size (*p >* 0.05), although within-subject trends remained consistently positive. Specifically, p-values were: Awake (*p* = 0.1514), N2 (*p* = 0.0678), N3 (*p* = 0.0539), and REM (*p* = 0.0128), indicating the most robust improvement in the REM stage. As expected, recall slightly decreased due to the stricter triggering criteria, particularly under the hybrid logic, and the latency introduced by the smoothing process. Notably, N1 classification remained poor across all methods, often reaching 0.0 recall, which is attributed to the low weight assigned to N1 in the model architecture. This can be addressed by future work, adjusting class weighting strategies.

**Table 4.** Comparison of Test Set Performance Across Control Strategies: Precision, Recall, and Trigger Count Before and After Smoothing (N=4)

|  | Threshold Cutoff |  | Naïve Waiting |  | Entropy Constraint |  |
| --- | --- | --- | --- | --- | --- | --- |
|  | Before | After | Before | After | Before | After |
| Awake | 0.663±0.220 | **0.744±0.176** | 0.624±0.254 | **0.663±0.220** | 0.667±0.215 | **0.795±0.166** |
| N2 | 0.931±0.034 | **0.952±0.022** | 0.909±0.035 | **0.931±0.034** | 0.932±0.036 | **0.955±0.017** |
| N3 | 0.959±0.022 | **0.981±0.019** | 0.938±0.035 | **0.959±0.022** | 0.945±0.033 | **0.959±0.025** |
| REM | 0.724±0.107 | **0.811±0.113** | 0.724±0.125 | **0.724±0.107** | 0.761±0.103 | **0.861±0.097** |

*Recall (N = 4, Mean ± SD)*
|  | Threshold Cutoff |  | Naïve Waiting |  | Entropy Constraint |  |
| --- | --- | --- | --- | --- | --- | --- |
|  | Before | After | Before | After | Before | After |
| Awake | **0.953±0.021** | 0.845±0.060 | 0.932±0.049 | **0.953±0.021** | **0.914±0.030** | 0.808±0.080 |
| N2 | **0.830±0.049** | 0.808±0.065 | **0.905±0.060** | 0.830±0.049 | **0.826±0.058** | 0.770±0.082 |
| N3 | **0.912±0.032** | 0.883±0.034 | **0.931±0.040** | 0.912±0.032 | **0.943±0.034** | 0.919±0.046 |
| REM | **0.758±0.218** | 0.658±0.295 | **0.841±0.171** | 0.758±0.218 | **0.696±0.242** | 0.602±0.312 |

*# of Trigger (N = 4, Mean ± SD)*

|  | Threshold Cutoff |  | Naïve Waiting |  | Entropy Constraint |  |
| --- | --- | --- | --- | --- | --- | --- |
|  | Before | After | Before | After | Before | After |
| Awake | **168.5±59.726** | 139.5±67.884 | **178.3±56.751** | 168.5±59.726 | **160.8±58.789** | 129.8±69.611 |
| N2 | **331.5±111.2** | 317.500±112.0 | **370.5±124.797** | 331.5±111.161 | **330.0±112.377** | 304.0±115.336 |
| N3 | **163.5±24.047** | 155.0±24.104 | **170.8±24.944** | 163.5±24.04 | **172.0±26.693** | 164.8±24.014 |
| REM | **106.3±23.541** | 79.5±34.158 | **121.0±21.749** | 106.3±23.51 | **92.0±28.766** | 67.0±36.104 |

*\*Note:* In this table, **Before** refers to the result of applying the specified control logic (indicated in bold above) to the raw output of the model. **After** refers to the result of applying the same control logic to the smoothed output. Importantly, this table was constructed to prove the hypothesis that applying smoothing *Before* control logic may yield higher precision than applying control logic directly to the raw output. The unmodified raw output from the ANN model (without any control logic) yields the following average precision scores across $N = 4$ test subjects (*Mean ± SD*): Awake = $0.605 \pm 0.256$ , N2 = $0.910 \pm 0.041$ , N3 = $0.930 \pm 0.043$ , REM = $0.660 \pm 0.124$ .

Despite a small reduction in recall, the number of valid triggers remained clinically adequate in smoothed conditions. For example, under N2-targeted stimulation, the average number of smoothed triggers per subject remained practical: Wake (129.8 ± 69.6), N2 (304.0 ± 115.3), N3 (164.8 ± 24.0), and REM (67.0 ± 36.1); all within acceptable operational ranges for closed-loop stimulation paradigms. To further boost precision, we analyzed a hybrid logic, combining threshold (*>* 0.7) and entropy (*<* 1.0) constraints using a logical AND operation. This strategy yielded the highest overall precision while maintaining a manageable trigger count (Table 4, Table 5). The visualization in Fig. 9 illustrates this multi-stage pipeline: smoothing followed by a hybrid constraint substantially improves alignment with ground-truth hypnograms for Subject 2. Panels C and D in Fig. 9 compare predicted hypnograms before and after smoothing, showing clear reductions in oscillatory misclassifications. Panel E shows the N2-targeted trigger events (red lines) overlaid on expert-annotated staging, confirming that most stimulations now align with true N2 windows. Panels F and G zoom in on the first 2 hours, showing corresponding EEG time-frequency maps and raw EEG traces, further validating the effectiveness of the smoothing and hybrid-triggering pipeline. Together, these results demonstrate that the proposed pipeline effectively reduces noisy activations and improves real-time precision for safe and reliable neuromodulation.

**Table 5.**
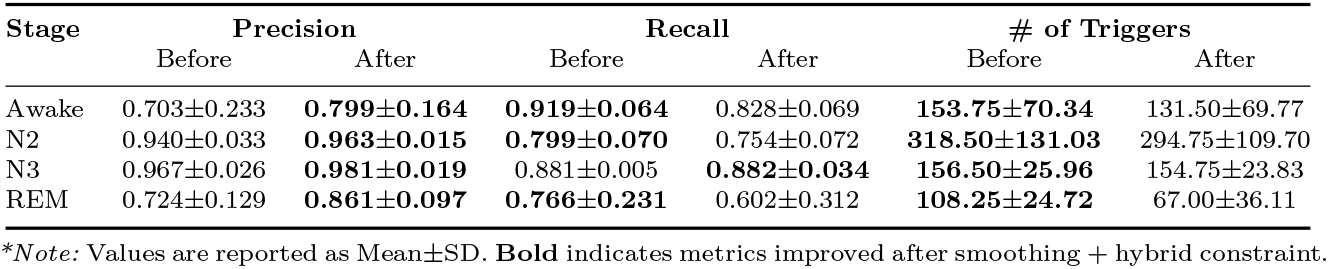
Performance of Hybrid Constraint Strategy Before and After Smoothing. Precision, Recall, Trigger Count (N = 4, Mean ± SD)

| Stage | Precision |  | Recall |  | # of Triggers |  |
| --- | --- | --- | --- | --- | --- | --- |
|  | Before | After | Before | After | Before | After |
| Awake | 0.703±0.233 | **0.799±0.164** | **0.919±0.064** | 0.828±0.069 | **153.75±70.34** | 131.50±69.77 |
| N2 | 0.940±0.033 | **0.963±0.015** | **0.799±0.070** | 0.754±0.072 | **318.50±131.03** | 294.75±109.70 |
| N3 | 0.967±0.026 | **0.981±0.019** | 0.881±0.005 | **0.882±0.034** | **156.50±25.96** | 154.75±23.83 |
| REM | 0.724±0.129 | **0.861±0.097** | **0.766±0.231** | 0.602±0.312 | **108.25±24.72** | 67.00±36.11 |

*\*Note:* Values are reported as Mean±SD. **Bold** indicates metrics improved after smoothing + hybrid constraint.

**Figure 9.**
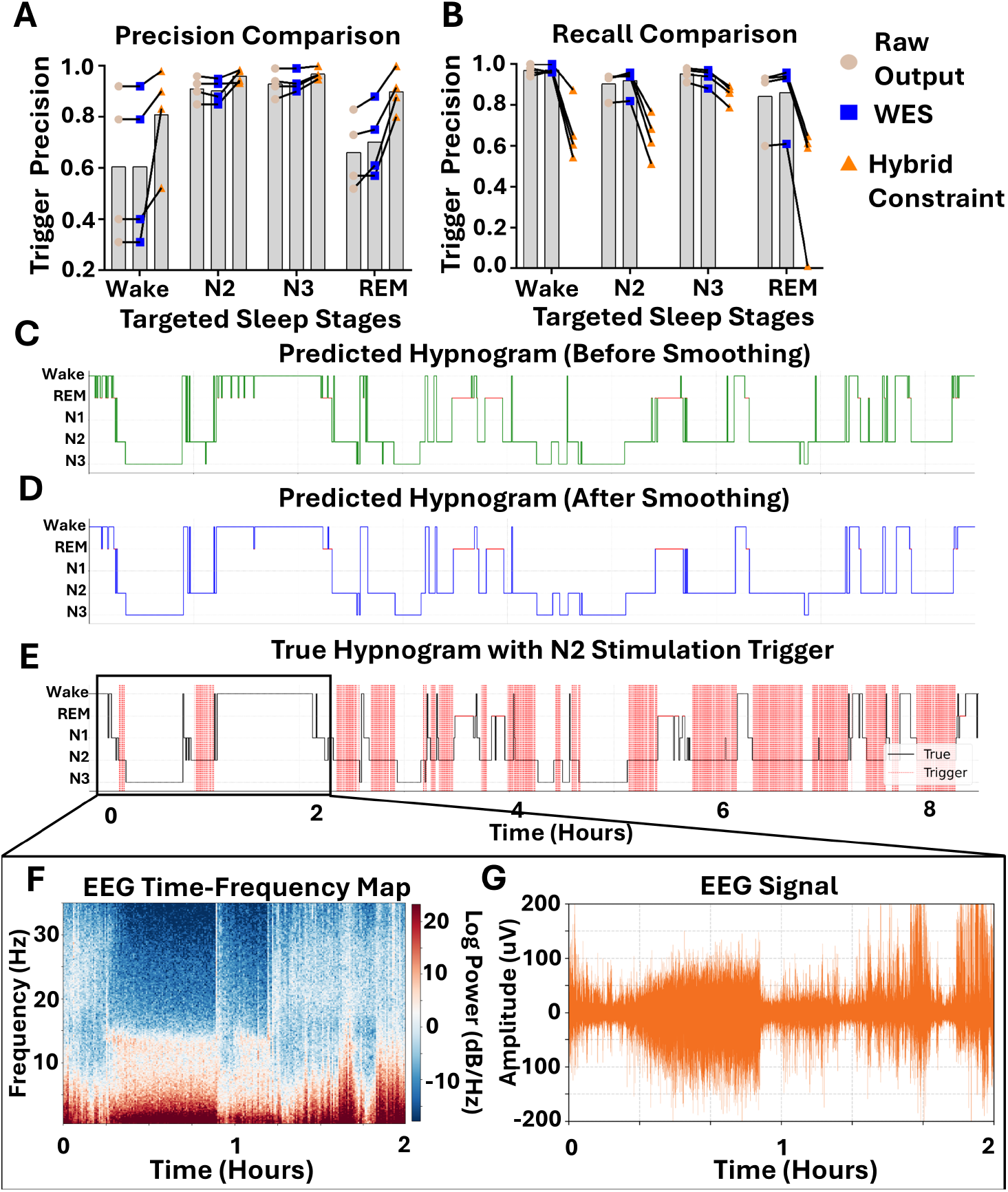
Progress of Trigger Precision and Recall through Smoothing and Hybrid Constraint Logic, with Detailed Visualization of N2-targeted Stimulation Alignment in a Representative Subject. Sleep-stage triggers were evaluated across four test recordings at three stages of the post-processing pipeline: raw model output, WES smoothing, and hybrid constraint logic (entropy *<* 1.0 & probability *>* 0.7). **(A, B)** Trigger *precision* (A) and *recall* (B) for Wake, N2, N3, and REM across the three pipeline stages: raw model output (beige), WES smoothing (blue), and hybrid constraint logic (orange). N1 was excluded due to persistent classification failure across all methods. Hybrid logic consistently yields the highest precision while introducing a trade-off with recall. Each connected line indicates the same subject. **(C, D)** Predicted hypnogram of test Subject 2 before (C, green) and after (D, blue) WES smoothing. Oscillatory misclassifications and short false REM segments (red highlights) are notably reduced. The effect was especially noticeable approximately 2 hours after the start (reduced REM ↔ Awake oscillation). **(E)** Ground truth hypnogram (black) with N2-targeted stimulation triggers (red lines) generated using hybrid constraint logic, demonstrating alignment with true N2 epochs. A detailed view of the first two hours is shown in panels (F) and (G). **(F)** Time–frequency representation (0-35 Hz, log power) of FPz EEG over the first 2 hours for Subject 2, showing clear separation of sleep stages, aligned with true hypnogram. **(G)** Raw FPz EEG amplitude trace over the same time segment, confirming electrophysiological validity of detected N2 epochs and stimulation alignment.

### 3.6. Transition Presence Matrix and Biological Plausibility

While ES achieved slightly better average performance across subjects, we evaluated the proposed WES as a non-recursive alternative. Unlike ES, which recursively blends current predictions with a single prior smoothed output, WES explicitly accumulates evidence from multiple past predictions over a longer fixed-length window using decaying weights. This design enabled integration with a transition prior matrix derived from empirical sleep dynamics. We also evaluated the Transition-Constrained WES (TC-WES), which modulated past predictions using a normalized transition matrix. While the average performance gain of TC-WES over WES was modest (*<* 1%), one subject exhibited *>* 5% improvement, suggesting possible benefits in specific cases.

To formalize biologically plausible transitions, we constructed a binary transition presence matrix for each of the 29 subjects based on observed stage-to-stage changes. These matrices were then averaged and normalized to generate a subject-agnostic probabilistic prior, representing the empirical likelihood of each transition occurring (Fig. 10 The resulting matrix highlighted transitions that never occurred, such as REM → N3 or Wake → N3, as biologically implausible. This matrix identifies transitions that never occurred in the dataset (e.g., Wake → N3) and was used to constrain the TC-WES model. (*Note:* we did a smooth penalization of probability, not a rejection of impossible transitions). The corresponding directed graph (Fig. 10) reflects permitted transitions and was integrated as a transition constraint during smoothing. While this matrix does not capture frequency, it provides a foundation for future studies to incorporate individualized or pathology-specific sleep dynamics, such as in insomnia patients or in populations with abnormal sleep architecture.

**Figure 10.**
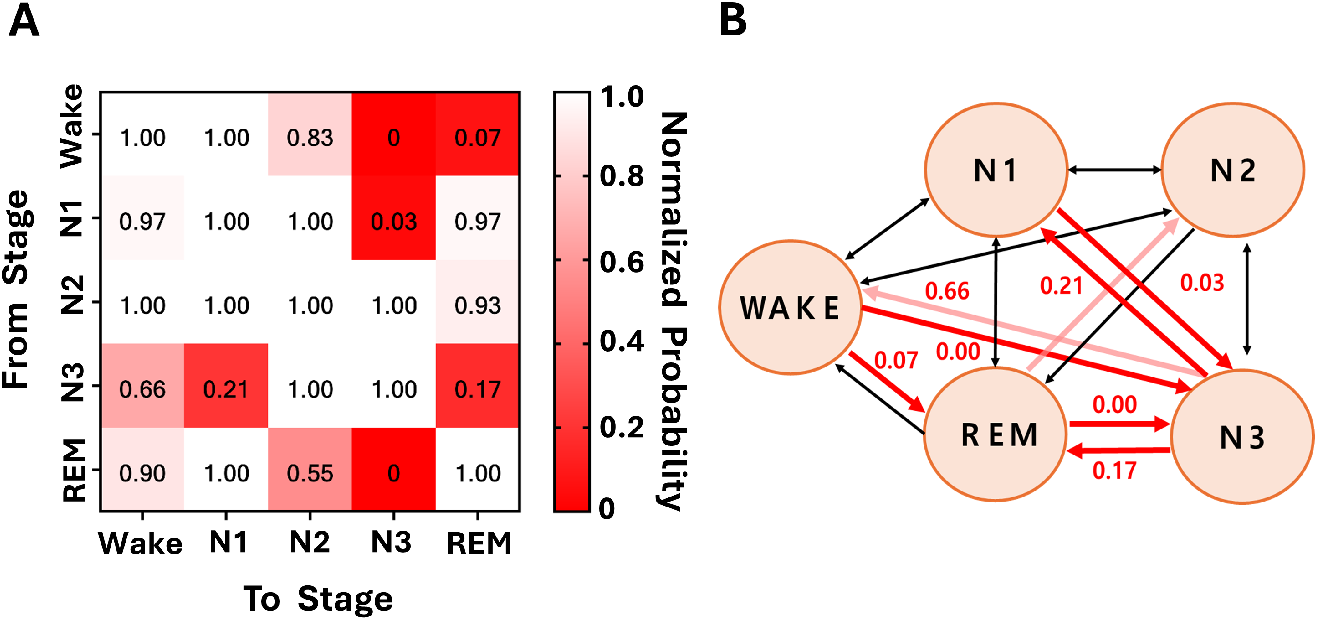
Transition Possibility Matrix and State Diagram Across 29 Subjects. **(A)** Binary transition presence matrix across all subjects (1: possible, 0: absent), where red indicates transitions that have scarcely occurred in any subject. Values represent the proportion of subjects in which each transition was observed. Highly improbable transitions include REM → N3 (0.00), Wake → N3 (0.00), N1 → N3 (0.03), Wake → REM (0.07), and N3 → REM (0.17). Moderately improbable transitions include N3 → N1 (0.21), REM → N2 (0.55), and N3 → Wake (0.66), collectively suggesting that certain abrupt or reverse transitions are biologically implausible and should be penalized or filtered in post-hoc correction or model constraints. **(B)** Corresponding directed graph of allowed transitions based on (A), used as a constraint in the TC-WES smoothing model. Edge weights and thickness reflect transition presence scores. *Note*: this matrix captures the existence —not frequency—of transitions. For example, value 1.0 from Stage Awake to Stage N3 explains that the transitions from Wake to N3 were observed across all subjects, and are not meant to always happen.

## 4. Discussion

This study presents a comprehensive, generalizable post-processing framework that enhances the reliability, precision, and real-time feasibility of sleep stage classification in closed-loop neuromodulation systems. Our findings offer several key contributions:

First, we demonstrate that temporal smoothing, particularly Weighted Exponential Smoothing (WES) and standard Exponential Smoothing (ES), significantly improves classification performance and robustness. Moderate smoothing yielded 1–2% absolute improvements in macro F1-score, ∼ 28.8% entropy reduction, and preserved low-latency responses (*<* 2.5 minutes). In high-performing models ( ∼ 8-90% baseline accuracy), such small gains are non-trivial and meaningful, as performance improvements typically plateau without costly increases in model complexity, training time, or dataset size. Therefore, these suggested smoothing parameters may help to reduce the need for an extensive optimization process, offering a practical means to improve stability without compromising latency or accuracy. Our lightweight, post-hoc solution instead achieves generalizable and interpretable improvements with minimal computational time(*<* 0.1 sec), making it ideal for real-time wearable systems.

While ES performed strongly overall, WES offers distinct conceptual and practical advantages. As a non-recursive evidence accumulation method, WES incorporates richer temporal context and allows for modular extensions, such as the application of physiological constraints or adaptive weighting, at a granular level. Unlike ES, which is sensitive to high smoothing parameters due to its recursive nature, WES distributes influence across past predictions in a larger window, enhancing robustness to noisy or unstable inputs. These properties make WES particularly suitable for closed-loop systems operating under unpredictable physiological and environmental conditions.

Second, we show that control logic applied after smoothing, particularly hybrid approaches combining high-confidence (probability threshold) and low-uncertainty (entropy constraints), can further increase stimulation precision. The hybrid logic consistently yielded high precision across stages (N3: 98%, N2: 96%, REM: 86%, Wake: 80%), suggesting its utility in stage-specific neuromodulation triggers, especially where false positives pose clinical risks.

Third, we explored a Transition-Constrained Weighted Exponential Smoothing (TC-WES) approach, designed to suppress biologically implausible transitions, such as Wake → REM, by softly penalizing low-probability stage changes during the smoothing process. This method extends prior work applying transition constraints in Hidden Markov Models (HMMs) to more general neural network-based models [30]. By embedding physiological priors directly into the inference pipeline, TC-WES enables more biologically grounded predictions without relying on rigid post-hoc rules.

While the overall improvement of TC-WES over standard WES was smaller than expected, it produced meaningful gains in certain individuals (e.g., *>* 5% improvement in one subject), suggesting potential for personalized or condition-specific adaptations. To understand the limited aggregate gains, we examined model behavior in detail. Our base classifier tended to increase REM probabilities gradually while maintaining relatively high and stable Wake probabilities over time. Because Wake → REM transitions are extremely rare in the transition matrix, the constraint mechanism disproportionately favored Wake, even when REM was the correct label. This led to suppression of true REM predictions and prolonged misclassification of Wake, inadvertently reinforcing the model’s existing bias.

Despite these challenges, TC-WES represents a novel avenue for integrating soft physiological constraints into real-time neural inference. Future work may improve its effectiveness by dynamically adjusting constraint strength, incorporating model-aware correction, or adapting to individual-specific sleep architectures.

Our work also identified practical considerations for real-world deployment. The proposed channel selection strategy used a minimal number of electrodes concentrated on one facial hemisphere, allowing full-brain coverage while enabling a feasible adhesive wearable design, an important step toward non-intrusive, home-based neuromodulation systems.

This study has several limitations. First, our dataset consisted of 29 healthy subjects with limited demographic variability, which constrains the statistical power and generalizability of our findings. Although we conducted subject-independent testing and performed robustness analyses using synthetic noise, these scenarios remain virtual approximations. Real-world environments introduce additional sources of interference, including stimulation artifacts, electromagnetic interference (EMI), and movement-related noise, that are not fully captured in our current simulations. Future work should validate the smoothing framework under these realistic conditions and across more diverse clinical populations

Second, our delay and mismatch analyses were conducted retrospectively by post-hoc alignment to ground truth transitions. This may exclude missed detections, misaligned predictions, and inflate or underestimate latency. Since true ground-truth correspondence is difficult to establish in noisy, dynamic systems, our reported delays should be viewed as conservative upper bounds. Moreover, because each 30-second EEG epoch is required before producing a prediction as a model input, a minimum latency of one epoch is inherent. However, this minimal latency could be reduced by decreasing the window size. In practice, real-time systems utilizing smoothing and control strategies are likely to trigger earlier than retrospective alignment suggests, often within 1-2 epochs ( ∼ 0.5-1 min).

To more precisely isolate the effect of smoothing on delay, we also compared raw output (not true label) versus smoothed outputs. While this approach avoids conflating smoothing delay with model reaction time, it assumes raw predictions are temporally accurate, which may not always hold under uncertainty or noise. Nonetheless, in most cases, observed delays remained under 1 minute, indicating that the added latency introduced by smoothing is practically minimal.

Several avenues for future work arise from these findings. First, integrating the proposed smoothing and control logic framework with state-of-the-art real-time sleep staging models (e.g., DeepSleepNet [27], DREAMT [63], or Transformer-based architectures [28]) could enable joint optimization of temporal dynamics and model predictions. Unlike recurrent models (e.g., LSTM [26]), which learn temporal dependencies within an epoch, smoothing operates across epochs, capturing longer-range consistency. Comparative studies could reveal how these different mechanisms, internal memory in sequence models versus external smoothing, affect robustness to noise, precision-recall tradeoffs, and temporal stability. In particular, applying smoothing to Transformer-based models may yield new insights regarding the relationship between multiple epochs. Transformers encode sequential dependencies via self-attention across the input window. Investigating how smoothing layers interact with attention-based memory (e.g., whether they complement or interfere with temporal continuity) could guide the design of more stable and interpretable sequence models for real-time applications.

Second, the transition presence matrix introduced here opens the path to individualized or disorder-specific constraints. Future versions may dynamically adapt the matrix over time or be used as a data-driven analog to sleep quality indices as well as the Pittsburgh Sleep Quality Index (PSQI), enabling automatic detection of abnormal transitions (e.g., Wake ↔ REM in narcolepsy or prolonged N1 in insomnia).

Third, while our work focuses on sleep neuromodulation, the smoothing + control logic framework generalizes to other noisy, sequential biosignal tasks. For example, maximizing precision may benefit seizure detection, muscle activation mapping, or dynamic safety triggers in implantable devices. In such domains, minimizing false positives and suppressing oscillatory misclassifications are critical for energy conservation (e.g., Pulse Width Modulation) and patient safety.

## 5. Conclusion

Taken together, our results establish a robust, interpretable, and lightweight framework for improving real-time sleep stage classification. By combining temporal smoothing with precision-enhancing control logic, we provide a generalizable method that not only improves performance under uncertainty but also aligns with clinical constraints for reliable, adaptive stimulation. This work lays the foundation for future development of physiologically-informed, closed-loop neuromodulation systems that extend well beyond sleep.

## Data Availability Statement

The data used in this study are publicly available as part of the Open Source Framework for Analyzing Physiology (OSF-ANPHY) sleep dataset, which comprises overnight high-density EEG polysomnography recordings from 29 healthy adults [37]. The custom code supporting the findings of this study is available from the corresponding author upon reasonable request.

## Conflict of Interest

The authors declare that there is no conflict of interest regarding the publication of this paper.

## Funding

Huiliang Wang would like to acknowledge support from the Department of Defense, Defense Advanced Research Projects Agency (DARPA) (REM-REST) grant (HR00112490328), Alzheimer’s Association New to the Field research grant (AARG-NTF-22-965140), Texas Proof of Concept (POC) Awards and Cockrell Innovation Grant. Tony Chae would like to acknowledge support from the James Sulzer Award for Undergraduate Research Fellowship at the 2025 9th Cellular to Clinically Applied Rehabilitation Research and Engineering (CARE) Symposium.

## Author Contribution

T.S.C led the project, developed the core methodology, performed experiments and analysis, and drafted the manuscript. J.d.R.M. and H.W. supervised the project. All other authors contributed through technical discussions, critical feedback, or revision suggestions.

## Acknowledgments

This manuscript was partially refined using ChatGPT-4o (OpenAI) for grammar, translation, and formatting assistance. The authors would like to thank to Dr. Ju-Chun (Leon) Hsieh for providing valuable technical advice.

## Notes

### Competing Interest Statement

The authors have declared no competing interest.

